# Actin dynamics sustains spatial gradients of membrane tension in adherent cells

**DOI:** 10.1101/2024.07.15.603517

**Authors:** Juan Manuel García-Arcos, Amine Mehidi, Julissa Sánchez Velázquez, Pau Guillamat, Caterina Tomba, Laura Houzet, Laura Capolupo, Giovanni D’Angelo, Adai Colom, Elizabeth Hinde, Charlotte Aumeier, Aurélien Roux

**Affiliations:** Department of Biochemistry, University of Geneva, CH-1211 Geneva, Switzerland; School of Physics, University of Melbourne, Parkville, VIC 3010, Australia; School of Life Sciences, Ecole Polytechnique Fédérale de Lausanne, CH-1015 Lausanne, Switzerland; Biofisika Institute (CSIC, UPV/EHU) and Department of Biochemistry and Molecular Biology, University of the Basque Country, ES-48940 Leioa, Spain; Ikerbasque, Basque Foundation for Science, ES-48013 Bilbao, Spain; Department of Biochemistry and Pharmacology, University of Melbourne, Parkville, VIC 3010, Australia

**Keywords:** Membrane tension, cell mechanics, actin dynamics, actomyosin, cytoskeleton, cell migration, biophysics

## Abstract

Tension propagates in lipid bilayers over hundreds of microns within milliseconds, precluding the formation of tension gradients. Nevertheless, plasma membrane tension gradients have been evidenced in migrating cells and along axons. Here, using a fluorescent membrane tension probe, we show that membrane tension gradients exist in all adherent cells, whether they migrate or not. Non-adhering cells do not display tension gradients. We further show that branched actin increases tension, while membrane-to-cortex attachments facilitate its propagation. Tension is the lowest at the edge of adhesion sites and highest at protrusions, setting the boundaries of the tension gradients. By providing a quantitative and mechanistic basis behind the organization of membrane tension gradients, our work explains how they are actively sustained in adherent cells.

## Introduction

Cells and organelles are separated from their environment by lipid membranes, which are self-healing fluid films,^1^ conferring them with unique mechanical properties. The main modulator of membrane mechanics is membrane tension, defined as the stress resulting from changing the apparent surface area of the membrane.^2^ In cells, plasma membrane tension coordinates endo- and exocytosis rates both temporally and spatially,^3–6^ determines cell shape during single-cell migration,^7–13^ and collective migration,^14^ and regulates membrane signaling,^15,16^ actin polymerization,^17^ axon growth,^18,19^ and cell spreading.^20,21^

Among the open questions about cellular membrane mechanics, the propagation of membrane tension in the plasma membrane is still under debate.^22^ Simply put, if membrane tension propagates too fast, tension gradients cannot arise. In pure lipid membranes (i.e.: giant unilamellar vesicles, GUVs), because of their fluidity, tension is expected to propagate at the speed of sound waves^23^ (>0.1 m/s), causing equilibration within tens of milliseconds and preventing observable tension gradients upon local perturbations.^24^ However, membrane tension gradients have been observed in supported lipid bilayers adhering to a solid substrate during expansion, which dramatically reduces the diffusion of lipids.^1,25^ In cells, early studies of the membrane tension of axons showed that the membrane flowed from the growth cone to the soma driven by a membrane tension gradient.^3^ While this gradient could be specific to the unique strong membrane-cytoskeleton interactions of axons,^26–29^ more recent studies have established that similar gradients can spread from the front to the rear of migrating cells.^8,12^ During cell migration, tension propagation is considered to be fast (< 1s across the cell) and long-ranged (at the cellular scale) and was proposed to act as a global signal transducer.^30^ But the maintenance of the gradient in this case is proposed to be linked to the high dynamics of actin forces. The current understanding is that the propagation of tensile stresses throughout the cell membrane depends on the cell type and the kind of perturbation, but that at rest, no cell exhibits tension gradients.^19,31–35^ In analogy to *in vitro* experiments, these divergences might originate from strong mechanical interactions between the membrane and the underlying actomyosin cortex or the substrate.^31,36,37^ However, whether these gradients can be generated in every cell is unknown, and a formal quantitative assessment is required to establish how they are formed.

Another layer of complexity in cellular membranes is their lateral separation into nanodomains.^38–43^ The lateral heterogeneity of cell membranes stems from a complex lipid composition constantly stirred by the dynamics of the underlying actomyosin cortex.^44,45^ Domain formation depends on temperature and lipid composition, but mechanical factors can play a role as well: studies *in vitro* have shown that, around a critical point,^46^ adhesion^47^ or increasing membrane tension by hypotonic shocks drives lateral phase separation.^48–53^ Interestingly, in model membranes far from critical points, increasing tension leads to an increased lipid spacing (less packing), as reported by Laurdan^54,55^ or Flipper-TR probes.^24^ In cells, results suggest membranes are near a critical transition point:^46^ increasing plasma membrane tension by hypotonic shocks globally promotes domain formation^56^ and increased lipid packing reported with Flipper-TR^24^ and Laurdan membrane probes.^57^ This link between membrane mechanics and molecular organization allows for proxy measurements of membrane tension using lipid packing-sensitive probes such as Flipper-TR, by determining the probe conformation by fluorescence lifetime imaging microscopy (FLIM).^24,58^ Importantly, the relation between lipid packing and tension does not depend on whether the experiments are performed with model membranes or in cells, but on the critical state of the membrane.

Membrane tension gradients were evidenced by differences in the force required to hold plasma membrane tubes (tether force) pulled from different parts of the cell. Tether force includes contributions not only from plasma membrane tension but also from membrane-cytoskeleton interactions and bilayer bending rigidity,^37,59–63^ which is composition-dependent.^64^ Therefore it remains unclear to what extent the reported changes in tether force stem from actual membrane tension gradients, from changes in membrane-cortex binding energy, or from lipid composition changes. Moreover, limited by discrete spatial measurements of the dorsal cellular plasma membrane, tube pulling cannot provide insights into the shape of the gradients nor investigate the tension distributions of the ventral cell membrane in contact with the substrate. To visualize membrane tension gradients in cells and investigate the mechanisms that could sustain them, we aimed at using the Flipper-TR probe.

## Results

### Flipper-TR reports membrane tension gradients in reconstituted membranes

While Flipper-TR lifetime was shown to report changes in membrane tension, we wondered if it could report membrane tension gradients. To test whether Flipper-TR can report membrane tension gradients, we reconstituted tension gradients in model lipid membranes in vitro, consisting of expanding glass-supported lipid bilayers (SLB) with well-controlled lipid composition and stable tension gradients, as previously reported.^1,25^ In brief, lipids were dried over large silica beads acting as lipid carriers and placed over plasma-cleaned glass. Upon hydration, lipids wet the clean glass and expand from the bead outwards as a single bilayer. SLB expansion is counteracted by the friction of the membrane onto the glass, allowing the formation of a tension gradient (Figure 1A).^1^ SLBs expand rapidly at first and slow down following a power-law while the tension gradient relaxes (Figure S1A),^1,25^ allowing us to sample a range of tension gradients (see Methods).

**Figure 1:**
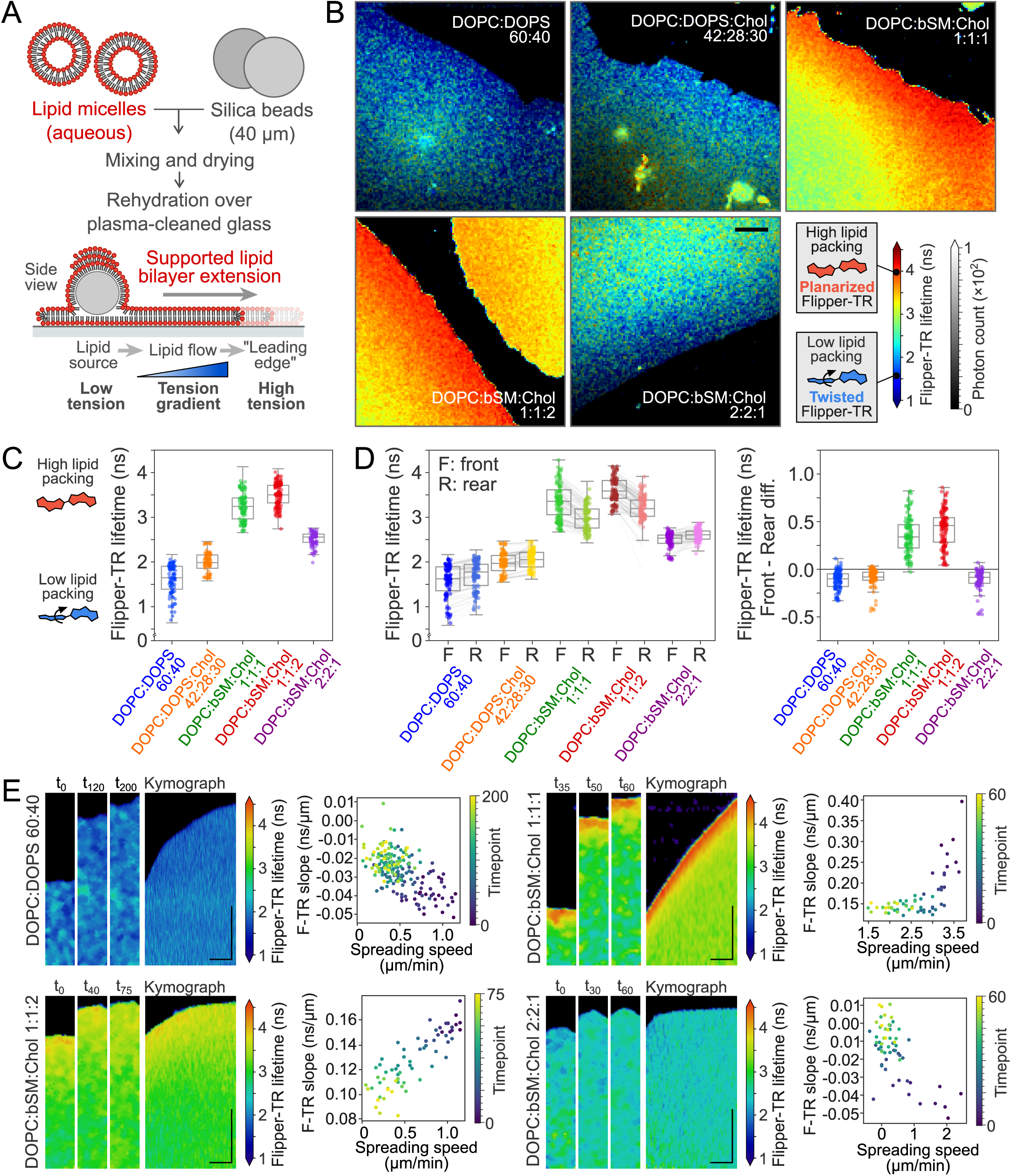
Flipper-TR reports membrane tension gradients in reconstituted membranes. **A**: Schematics of supported lipid bilayer experiments. **B**: Examples of expanding supported lipid bilayers with different compositions. Color indicates average Flipper-TR fluorescence lifetime (ns); black mask shows average photon count. **C**: Average Flipper-TR fluorescence lifetime in supported lipid bilayers with different compositions, spatially averaged over the field of view. **D**: Front (expanding edge) and rear (closest to lipid source) average Flipper-TR fluorescence lifetime in supported lipid bilayers with different compositions. Lines represent data pairing. Right, front-rear differences for different lipid compositions. **E**: Left, representative kymographs of expanding supported lipid bilayers of different compositions. Right, linear fits of spatial Flipper-TR gradients (ns/µm) over time. **B,E**: Scale bar, 10 µm.

We performed confocal FLIM of the SLBs with Flipper-TR using 5 different lipid compositions containing Dioleoylphosphatidylcholine/serine (DOPC/S), brain sphingomyelin (bSM), and cholesterol (Chol): DOPC:DOPS 60:40; DOPC:DOPS:Chol 42:28:30; DOPC:bSM:Chol 1:1:1, DOPC:bSM:Chol 1:1:2, and DOPC:bSM:Chol 2:2:1. Flipper-TR lifetime distributions qualitatively showed pronounced spatial gradients (Figure 1B), particularly evident in fast SLBs spreading at velocities comparable to cell migration (>5μm/min) (Supplementary Movie 2). As expected, more ordered membranes led to higher Flipper-TR lifetimes (Figure 1C). Interestingly, single-phase compositions display a lower average Flipper-TR lifetime at the front (leading edge) than the rear, closer to the lipid source, while some compositions containing phase-separating lipids (bSM and Chol) display an inverted lifetime gradient (Figure 1D). Thus, considering the gradient from the inner to the outer part of the membrane patch, compositions around a critical point display a positive slope while others display a negative slope.

Importantly, only bilayers that are expanding – thereby having a tension gradient – display a lifetime gradient (Figure S1A). To better understand the interplay between Flipper-TR lifetime and membrane tension, we plotted the slope of gradients as a function of the SLB spreading speed. For long time points, the speed reduces to zero as well as the slope value, confirming that when SLBs stop spreading and dissipate their tension gradient, the Flipper-TR lifetime gradient also vanishes (Supplementary Movie 2). For SLBs with compositions that cannot phase-separate, the slope of the Flipper-TR lifetime gradient (see Methods) increased from negative values to zero roughly proportionally to expansion speed (Figure 1E, DOPC:DOPS 60:40 and DOPC:bSM:Chol 2:2:1). For SLBs with compositions close to phase separation, the slope decreased instead from positive values to zero, also roughly proportionally to expansion speed (Figure 1E, DOPC:bSM:Chol 1:1:1 and DOPC:bSM:Chol 1:1:2). This effect is robust across a wide range of spreading speeds and lifetime gradients (Figure S1A).

Raedler and colleagues described that tracer fluorescent molecules display an exponentially decaying concentration profile in spreading SLBs with a tension gradient^1,25^. The decay length depends on the bulkiness of the tracer relative to the other lipids of the mix, probes with bulky headgroups such as DOPE-atto647 localize at areas with lower lipid packing, a visual evidence of membrane tension gradients.^1^ DOPE-atto647 localized to the edge of the spreading SLBs in single-phase lipid composition, but it was excluded from the edge of SLBs composed of phase-separating composition (Figure 1SB). This confirmed that in spreading SLBs under tension made of phase-separating compositions, a more ordered phase was colocalized with areas of higher tension.

Flipper-TR distribution also displayed fluorescence intensity profiles, inverted compared to the DOPE-A647 profiles (Figure 1SB). It is however more difficult to interpret these profiles in terms of concentration since the quantum efficiency of Flipper-TR varies with lipid order.^65^ However, the higher Flipper-TR fluorescence intensity at the edge of SLBs with phase-separating compositions is consistent with the presence of more ordered domains in these regions. Overall, these data showed the correspondence between tension and lipid packing and confirmed that Flipper-TR can report tension gradients in model membranes.

### Leading-edge extension increases Flipper-TR lifetime in migrating cells

Membrane tension gradients in migrating cells were first investigated in the fish keratocyte.^12^ Fish keratocytes have a very characteristic crescent shape and perform fast and persistent migration driven by a large lamellipodium at the leading edge.^10^ Knowing that Flipper-TR can report tension gradients, we wondered if we could visualize a lifetime gradient during keratocyte migration using Flipper-TR. Keratocytes were labeled with Flipper-TR following previously reported procedures^66^ and their ventral plasma membrane was imaged using confocal FLIM (see Methods). Our results confirmed that cells robustly display large lifetime differences between the lamellipodium leading edge and the cell rear (Figure 2A): ⟨Lifetime_front_⟩ = 4.85±0.13ns; ⟨Lifetime_rear_⟩ = 4.65±0.16ns; ⟨ΔLifetime⟩ = 0.20±0.12ns. This is consistent with the literature, reporting higher lipid packing and higher tension values at the front of migrating cells.^12^ At the level of single cells, front and rear values fluctuate over time, but the single-cell time average displays a robust gradient in Flipper-TR lifetime (Figure 2B).

**Figure 2:**
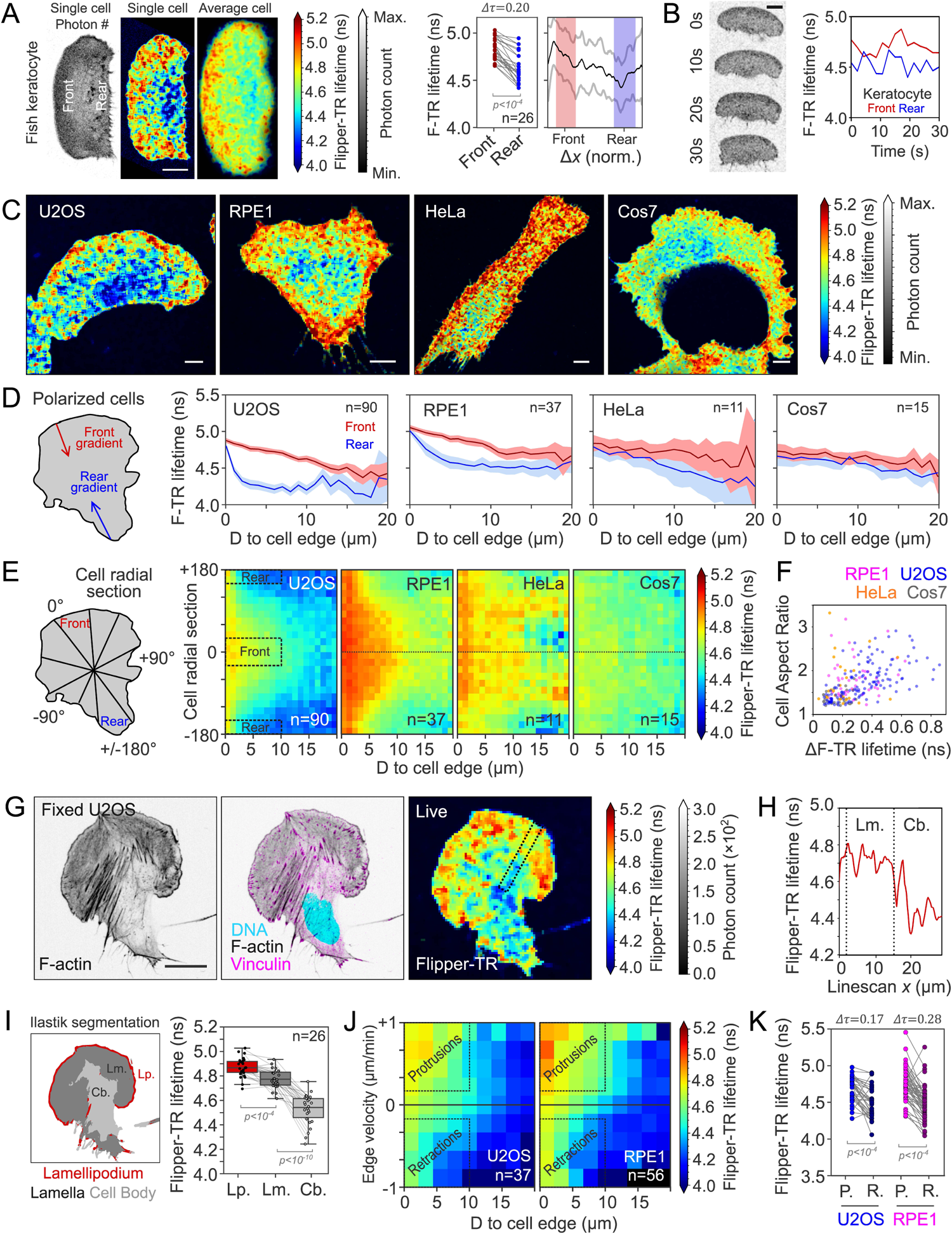
Leading-edge extension increases Flipper-TR lifetime in migrating cells. **A**: Left, representative live confocal FLIM microscopy image of fish keratocytes labeled with Flipper-TR. Color indicates average Flipper-TR fluorescence lifetime (ns); black mask shows average photon count. Right, average Flipper-TR fluorescence lifetime at the front and rear 20% areas of migrating fish keratocytes (n=27, Welch’s P<10^-4^), with lifetime profile along the normalized position (mean ± standard deviation). Front and rear regions are shaded in red and blue respectively. **B**: Left, representative time-lapse image of migrating fish keratocytes. Right, average Flipper-TR fluorescence lifetime at the front (red) and rear (blue) 20% areas during the time lapse. **C**: Representative image of polarized U2OS, RPE1, HeLa, and Cos7 cells labeled with Flipper-TR. Color indicates average Flipper-TR fluorescence lifetime (ns); black mask shows average photon count. **D**: Average Flipper-TR fluorescence lifetime as a function of distance from the edge at the front (red) and rear (blue) of polarized U2OS (n=90), RPE1 (n=37), HeLa (n=11), and Cos7 (n=15) cells. Line represents mean ± standard deviation. **E**: Average Flipper-TR fluorescence lifetime as a function of distance from the edge and radial position (front at 0°) of polarized U2OS (n=90), RPE1 (n=37), HeLa (n=11), and Cos7 (n=15) cells. Color indicates average Flipper-TR fluorescence lifetime (ns). **F**: Difference of average Flipper-TR fluorescence lifetime between front and rear 10-µm regions as a function of cell aspect ratio for U2OS (blue, n=132), RPE1 (magenta, n=43), HeLa (orange, n=63), and Cos7 (grey, n=32) cells. **G**: Left, representative image of fixed U2OS cells labeled with phalloidin-Alexa647, Hoechst, and anti-vinculin-alexa488. Right, live image of U2OS labeled with Flipper-TR. Dotted region indicates line scan region for panel H. **H**: Average Flipper-TR fluorescence lifetime along a line scan marked in panel G. Dotted lines indicate lamellipodium, lamella (Lm.), and cell body (Cb.). **I**: Left, segmentation of the representative cell in panel G into lamellipodium (red, Lp.), lamella (dark grey, Lm.), and cell body (light grey, Cb.). Right, average Flipper-TR fluorescence lifetime in these regions (n=26, paired Welch’s P). **J**: Average Flipper-TR fluorescence lifetime as a function of distance from the edge and edge velocity in U2OS (n=37), RPE1 (n=56) cells. Color indicates average Flipper-TR fluorescence lifetime (ns). **K**: Average Flipper-TR fluorescence lifetime at protruding (P) and retracting (R) regions in U2OS (n=37), RPE1 (n=56) cells (paired Welch’s P). **A,B,C,G**: Scale bar, 10 µm.

To extend these observations, different cell lines were selected to cover a wide range of migratory behaviors: U2OS and RPE1, which migrate efficiently, and HeLa and Cos7 which appear slow and less persistent (Figure S2A-B). Qualitatively, as for keratocytes, the cells showed large heterogeneities of Flipper-TR lifetime (Figure 2C), being higher at cell edges and decaying toward the cell center. Contrary to keratocytes, these cell lines do not maintain shape during migration, allowing us to manually sort cells into polarized and non-polarized categories based on their shape. While cells displaying a highly polarized shape showed clear differences between front and rear spatial decays (Figure 2C-D), non-polarized cells did not (Figure S2C-D). Lifetime values decayed linearly over more than 10 microns from the cell leading edge, while they decayed exponentially within a few microns to the baseline levels from the rear (Figure 2D-F). Lifetime differences between front and rear were more pronounced in cells with elongated morphologies (Figure 2F). Overall, the cell lines that displayed a more persistent migration (Figure S2A-B) were the ones that displayed the most robust Flipper-TR gradients. Interestingly, U2OS cells displayed overall lower lifetime values than other cell types but had the same gradient as the other persistent cell type RPE1. Since plasma membrane lipidomes are cell-type specific, offsets in the average lifetime may be due to a specific lipid composition in this cell line.^67^

We next investigated the mechanism leading to the observed differences in Flipper-TR lifetime. As the actin cortex shapes migrating cells, we wondered how the actin cytoskeleton would colocalize with Flipper-TR lifetime gradients. For this, following Flipper-TR lifetime imaging, U2OS cells were fixed and labeled for actin and vinculin (see Methods). In polarized cells, actin and vinculin had a stereotypical localization pattern composed of a dense, narrow actin strip marking the lamellipodium edge and a less dense actin region with small elongated focal adhesions corresponding to the lamella. At the rear, we observed low cortical actin and large stress fibers originating from large mature focal adhesions (Figure 2G). Regions of highest and lowest lifetime coincided with characteristic features in the actin organization. Flipper-TR lifetime increased both in the lamellipodium and lamella regions (Figure 2G, Flipper-TR panel, and Figure 2H). We used actin images to train a machine learning algorithm using Ilastik^68^ to segment cells into lamellipodium, lamella, and cell body regions, and calculate their average Flipper-TR lifetime values. This confirmed that the differences in actin cytoskeleton morphology led to different Flipper-TR lifetimes.

Actin polymerization extends the leading edge of lamellipodium and thereby could increase plasma membrane tension.^13,20,32,69^ To study this, we performed live FLIM on RPE1 and U2OS cells, to track cell edge dynamics simultaneously with Flipper-TR lifetime. FLIM time-lapses on migrating cells confirmed that, in the event of cell repolarization, the region of high lifetime dynamically changes localization to the new leading edge (Supplementary Movie 1). For every point of the cell edge, the gradient of Flipper-TR lifetime (from the cell edge inwards) and the edge velocity were measured in RPE1 and U2OS cells. Plotting the spatial lifetime gradient binned by edge velocity revealed that the protrusion speed is positively correlated with increased Flipper-TR lifetime (Figure 2J). The protruding parts of the cell display higher Flipper-TR lifetime values than the retracting ones (Figure 2K): ⟨U2OS Lifetime_protrusion_⟩ = 4.63±0.16ns; ⟨U2OS Lifetime_retraction_⟩ = 4.46±0.21ns; ⟨RPE1 Lifetime_protrusion_⟩ = 4.76±0.36ns; ⟨RPE1 Lifetime_retraction_⟩ = 4.47±0.22ns. These results are consistent with our hypothesis, thereby confirming that protrusions in migratory cells increase Flipper-TR lifetime.

Since the lowest lifetimes often correspond to the central region of the cells, coinciding with the nucleus position, we tested whether the presence of the nucleus could influence the reported plasma membrane Flipper-TR lifetime gradients. To test this, we characterized the lifetime of the ventral membrane of enucleated cells (Methods). Enucleated U2OS and RPE1 cells maintained similar Flipper-TR lifetime distributions (Figure S2E-F) supporting that the nucleus is not generating lifetime gradients within the plasma membrane.

### Membrane tension gradients are present in non-migrating cells

After confirming that Flipper-TR can reveal membrane tension gradients in expanding membranes both in migrating cells and *in vitro*, we tested if plasma tension gradients also exist in non-migrating cells. To this end, we cultured single HeLa cells on fibronectin-coated micropatterns^71,72^ and labeled them with Flipper-TR. Controlling cell shape allowed us to extract average lifetime maps from large sets of cells with the same shape, thus with comparable spatial organization of intracellular components, including the nuclei, the actin cytoskeleton, and the focal adhesions (Figure S3A). Flipper-TR lifetime showed a heterogeneous spatial distribution at the ventral plasma membrane (Figure 3A), as previously observed^5^, while its distribution appeared homogeneous on the dorsal membrane (Figure 3B). Despite no significant differences were observed in the average Flipper-TR lifetimes between different cell shapes, neither on the ventral/bottom (Figure 3A) nor dorsal/middle planes (Figure 3B), we found a significant shift in the average Flipper-TR lifetime value between the dorsal and ventral planes (Figure 3C, S3B): ⟨Lifetime_dorsal_⟩ = 4.98±0.05ns; ⟨Lifetime_ventral_⟩ = 4.71±0.06ns; such as ⟨ΔLifetime⟩ = 0.26±0.07ns.

**Figure 3:**
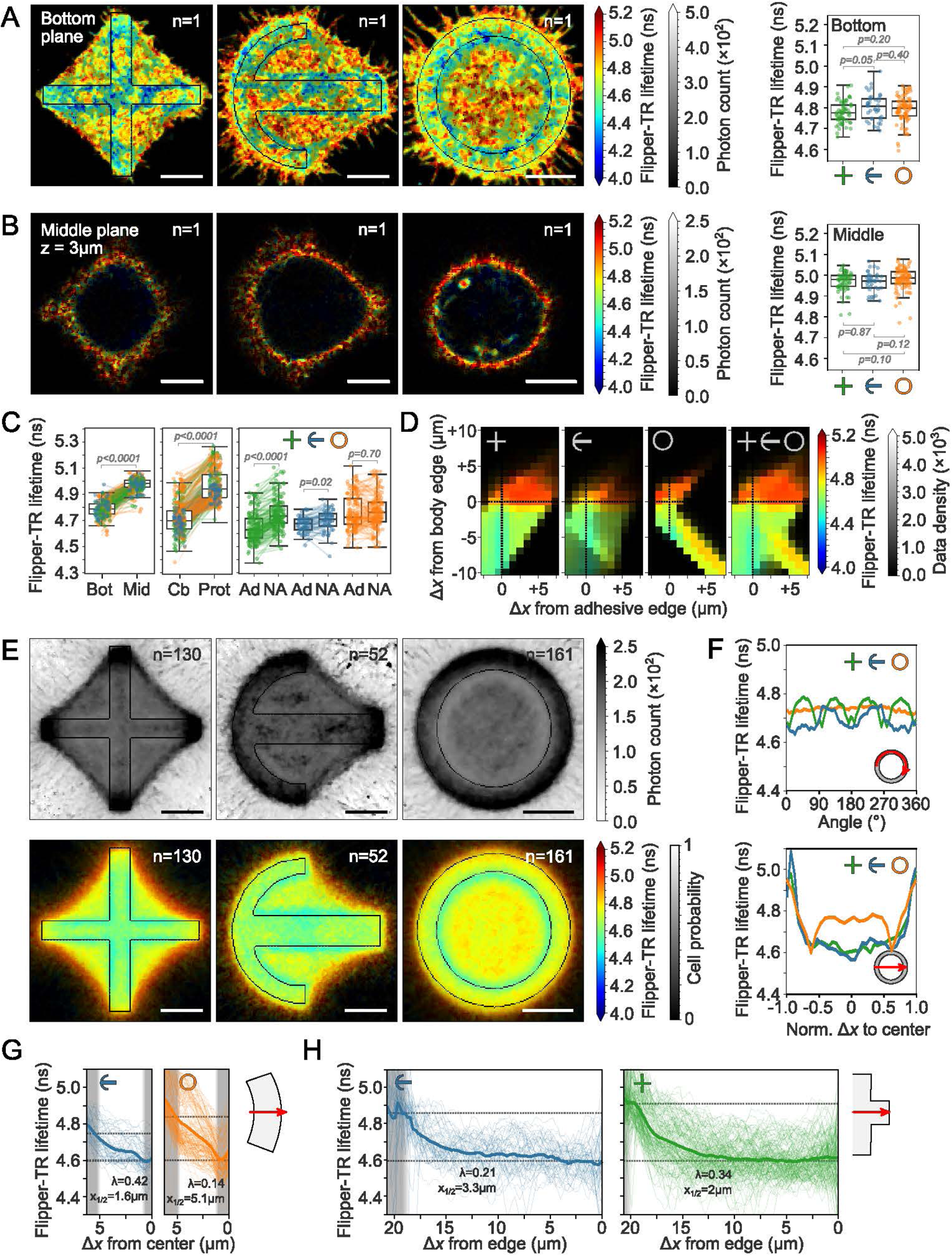
Tension gradients are also present in non-migrating cells. **A**: Left, representative images of ventral membrane (bottom plane) HeLa cells labeled with Flipper-TR on cross, crossbow, and ring micropatterns. Right, average Flipper-TR fluorescence lifetimes of cross, crossbow, and ring micropatterned cells (n=130, 52, 161, Welch’s P). Data distribution in black. **B**: Same as panel A, but on the dorsal plane (3 µm above the glass surface). **C**: Average Flipper-TR fluorescence lifetime (ns) in cell regions of cross (green), crossbow (blue), and ring (orange) micropatterned cells. Left, ventral (bottom) vs. dorsal (middle) planes. Center, cell body (Cb) vs. protrusive areas (Prot). Right, adhesive vs. non-adhesive areas of the cell body membrane. Data distribution in black. **D**: Average Flipper-TR fluorescence lifetime as a function of distance from micropattern edge and cell body edge. Color indicates average Flipper-TR fluorescence lifetime (ns); black mask shows data point density. **E**: Top, cell averaging of Flipper-TR photon counts in ventral membrane HeLa cells on micropatterns. Bottom, average Flipper-TR lifetime. **F**: Top, average Flipper-TR fluorescence lifetime along the cell contour of cross (green), crossbow (blue), and ring (orange) micropatterned cells. Bottom, lifetime distribution across the cell. **G**: Average Flipper-TR lifetime along pattern edges in crossbow (left, blue) and ring-patterned cells (orange). Shaded lines show individual traces. **H**: Average Flipper-TR lifetime across pattern in crossbow (left, blue) and cross-patterned cells (green). Shaded lines show individual traces. **G,H:** Grey bands represent non-patterned areas. **A,B,E**: Scale bar, 10 µm.

Consistent with the previous results, cellular membrane protrusions on the non-adherent areas exhibited higher lifetime values (Figure 3C). The average lifetime at the ventral membrane was comparable between adhesive and non-adhesive regions (Figure 3C). The spatial probability maps of Flipper-TR lifetime (Figure 3E) suggested that, while the gradients along the cell contour or across the cell diameter differ between shapes (Figure 3F), the average Flipper-TR lifetime increases at the cell edges and decreases over the boundaries of the adhesive micropatterns, regardless of shape. The lateral lifetime gradients between the regions of high and low lifetime were: ⟨Lifetime_high_⟩ = 4.85±0.13ns; ⟨Lifetime_low_⟩ = 4.67±0.13ns; such as ⟨ΔLifetime⟩ = 0.19±0.09ns. We further analyzed this by assigning two spatial descriptors to each lifetime value, namely the shortest distance cell edge and the shortest distance to the micropattern boundary. Note that negative distances account for pixels inside the cell or the micropattern. By plotting the average lifetime as a function of these two distances we can compare the lifetime gradients we obtained from the three different pattern shapes (Figure 3D). The fact that the gradients exhibited the same distribution regardless of the shape confirmed that the positions of the cell edge and the micropattern boundary are the main determinants of the Flipper-TR lifetime distribution (Figure 3G-H).

Epithelial RPE1 cells displayed similar spatial lifetime gradients decreasing from the cell edges to the cell center. Dorsal to ventral lifetime shifts were also conserved (Figure S3C-D). However, the increase of lifetime at plasma membrane regions over non-adhesive regions seems to be cell-line specific. This could be attributed to protrusive activity and ruffle formation over non-adhesive areas in HeLa (Supplementary movie 3), not present in RPE1. This was confirmed by visualizing average membrane tension gradients on different radial designs for HeLa cells, including full discs and double-ring patterns (Figure S3E). Cells on wider ring pattern lifetime decrease at the inner boundary of the pattern (Figure S3E). In full disc patterns, lifetime was distributed homogeneously (Figure S3E). Importantly, in all cases, Flipper-TR lifetime increased over non-adhesive regions in HeLa.

Despite the standardization of cell shape, cells still exhibit significant differences in vinculin and actin organization leading to two main protrusive behaviors: “low protrusion” cells display a very dense vinculin recruitment on focal adhesions and thick stress fibers and no protrusions; “high protrusion” cells display homogeneous vinculin localization and no stress fibers, with numerous protrusions projecting outwards (Figure S4A). By plotting the lifetime gradients of the 10% cells with the highest protrusion area and the 10% cells with the lowest we confirmed that cells with more protrusive behavior had higher lifetime values overall, suggesting, as in migrating cells, that actin protrusions play a fundamental role in the tension increase at the cells’ edges (Figure S4).

### Main lipid components are uniformly distributed in patterned cells

While our *in vitro* data showed that Flipper-TR can report membrane tension gradients in reconstituted membranes, we wondered whether the lifetime gradients we observed in cells had the same origin or if they could originate from lipid composition variations. To investigate this, we studied the spatial distribution of lipid composition by using Matrix-Assisted Laser Desorption Ionization Mass Spectrometry Imaging (MALDI-MSI). This technique allows the mass-spectrometry analysis of the chemical content of each pixel in an image providing a detailed analysis of most lipid species with a subcellular resolution.^73^ The subcellular resolution is achieved by averaging over many images, which was made possible using standardized cell shapes (Figure S3F and Methods).

We measured the fraction of different lipid species in cross-shaped micropatterned cells: namely phosphatidylcholine (PC), phosphatidylethanolamine (PE), ceramide (Cer), sphingomyelin (SM), and globotriaosylceramide (Gb3) (Figure S3G). From our results on model membranes, we expect that composition gradients can be linked to tension gradients only in lipids in trace amounts with extreme bulkiness (Figure S1B). Consistent with our hypothesis, lipid species that are abundant in these membranes (PC, PE) display homogeneous distributions while less abundant lipid species (Cer, SM, and Gb3) display heterogeneous spatial distributions in the form of lipid concentration gradients (Figure S3H). The spatial gradient of Gb3, which accumulates at the cell membrane protrusions is consistent with the tension gradient previously reported (Figure 3E), as tension is maximal at the cell edges (Figure S3H). Interestingly, the ceramide and sphingomyelin distribution showed a moderate preference for focal adhesion sites which do not correlate with an increase in Flipper-TR lifetime (Figure S3H). This supports that the Flipper-TR lifetime differences are not due to differences in lipid composition, as we could expect sphingolipids to increase membrane order. To further explore the spatial distribution of lipid composition throughout the plasma membrane, we used fluorescent dyes, Filipin, Cholera toxin, Equina toxin, and Shiga toxin that specifically bind to cholesterol, GM1, sphingomyelin, and Gb3 lipids, respectively (Figure S3I). Qualitatively, the concentration of all these lipid species tends to increase at the vertices of the cells on cross-shaped patterns, but the lipid packing effects for sphingolipids associated with lipid raft^74^ cannot explain the spatial variations observed in the average lifetime maps. Overall, these observations support that gradients reported by Flipper-TR lifetime are due to membrane tension gradients and not lipid composition.

### Actin polymerization and adhesions scale tension gradients

To test the role of actomyosin cytoskeleton in setting membrane tension gradients, we characterized the effect of several pharmacological treatments on the Flipper-TR lifetime gradients. We used 1µM Latrunculin A (‘LatA’) for inhibiting actin polymerization, 85 µM CK-666 for inhibiting Arp2/3-mediated actin nucleation, 10µM NSC668394 (‘NSC66’) for inhibiting ezrin – a major actin-membrane linker protein – and 20µM Y-27632 for inhibiting Rho kinase, responsible for myosin contractility. Moreover, we used the JLY cocktail consisting of 20µM Y-27632, 5µM Latrunculin B, and 8µM Jasplakinolide to block actin dynamics^75^. For each treatment and each shape, we quantified: i) the spatial lifetime distribution (Figure 4A top panels), ii) the radial lifetime distribution for the ring pattern (Figure 4A, bottom panels), iii) the average lifetime per cell (Figure 4B), and iv) the maximal amplitude of the gradients (Figure 4C).

**Figure 4:**
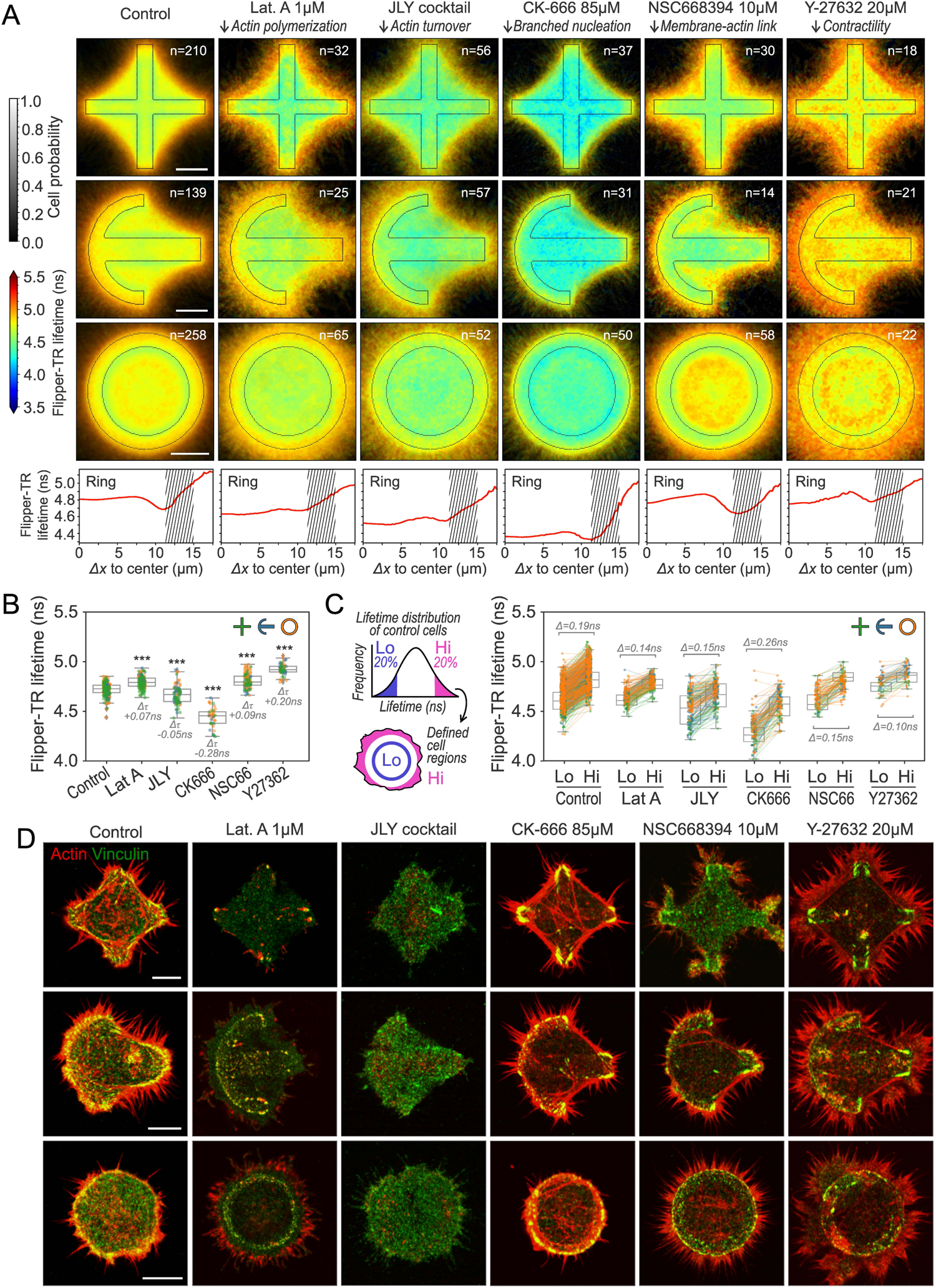
Actin polymerization and adhesions scale tension gradients. **A**: Top, cell averaging of average Flipper-TR lifetime in ventral membrane of HeLa cells on cross, crossbow, and ring micropatterns under different drug treatments. Bottom, average Flipper-TR lifetime vs. distance from center of ring micropattern. Shaded areas represent adhesive micropattern location. **B**: Average Flipper-TR lifetime (ns) per cell of cross (green), crossbow (blue), and ring (orange) micropatterned cells under different drug treatments. Labels represent Welch’s test P value (P<10^-3^) and difference from control in ns. Data distribution in black. **C**: Average Flipper-TR lifetime (ns) per cell of bottom and top quintiles (low/high) of cross (green), crossbow (blue), and ring (orange) micropatterned HeLa cells under different drug treatments. Labels display difference from control in ns. Lines represent data pairing. Data distribution in black. **D**: Representative fluorescence images at basal plane of cross, crossbow, and ring micropatterned cells stained with phalloidin (red) and vinculin antibody (green) under different drug treatments. **A,D**: Scale bar, 10 µm.

Both LatA and JLY treatments inhibited actin dynamics. As in control experiments, we observed a significant shift in the average lifetime between the ventral and dorsal membranes (JLY on Figures S4E-G). However, the increase in lifetime in non-adhesive areas and the pattern boundary effect were abolished. As a result, membrane tension became more homogeneously distributed at the ventral membrane, remaining high at the edges (Figure 4A and 4C). The linear decay of the Flipper-TR lifetime at the edge became exponential, decaying quickly towards the center. Therefore, LatA and JLY transformed front-like tension gradients to rear-like tension gradients through actin depolymerization. This suggests that the linear gradients observed in the lamellipodial regions of migrating cells (Figure 2) depend on the actin organization at the lamellipodium. F-actin was almost absent in both these conditions, although some focal adhesions remained in place (Figure 4D). These results confirm that actin dynamics is essential to generate membrane tension gradients.

In contrast with the LatA or JLY treatments, inhibiting the Arp2/3 complex with CK-666 did not decrease F-actin contents. Arp2/3 complexes nucleate about half of the cortical F-actin content in HeLa cells in the shape of branched networks.^76^ As expected, Arp2/3 inhibition led to the formation of more filopodia, larger lamellipodia, and thicker focal adhesions and stress fibers (Figure 4D). Surprisingly, the inhibition of Arp2/3 reduced considerably the average Flipper-TR lifetime value (Figure 4B). It nevertheless maintained both the higher values at the cell edges (Figure 4A and 4C) and the shift between the ventral and dorsal planes (Figure S4E). The lifetime drastically decreased at the lamellar regions, resulting in a steeper gradient than in control conditions (Figure 4A). This suggests that, in control conditions, lamellipodial extension – promoted by Arp2/3 branching – is generating higher membrane tension, and/or reducing the diffusion of lipids, setting a longer characteristic decay length of the gradient. The role of branched actin networks was also supported by Rho kinase inhibitor Y-27632 treatment, which significantly extended lamellipodia (Figure 4D). This resulted in a global increase in the average lifetime (Figure 4B), particularly in the numerous lamellipodia extending outwards (Figure 4A), and reduced the amplitude of the lifetime gradients (Figure 4C).

Finally, to inhibit membrane-to-cortex attachment, we used NSC668394, which inhibits ezrin activity by preventing its T567 phosphorylation, a member of the ERM protein family that acts as a major membrane-cortex linker.^77^ Cells under ezrin inhibition displayed similar protrusive morphology as the control cells but displayed smaller focal adhesions with less vinculin (Figure 4D). This resulted in a similar lifetime in ruffled areas, but a larger decay of lifetime in adhesive areas (Figure 4C). This indicates that with less membrane-to-cortex attachment can increase membrane tension at protrusions, but this tension can dissipate faster away from protrusive regions. We, therefore, concluded that actin-membrane attachment may be involved in reducing the local diffusion of membrane components, thereby reducing tension propagation and gradients. Overall, these results confirm that tension generation is controlled by the dynamics of the actin cytoskeleton, specifically of branched actin, via membrane-to-cortex attachment.

### Lipid diffusion and flows are limited in patterned cells

Previous studies have shown that gradients in membrane tension are associated with lipid flows,^18^ with lipids flowing from low to higher tension regions of neuronal growth cones. On the other hand, other studies on migrating cells have shown either no flow^78^ or rearward flow^79^ in the cell reference frame. We therefore wanted to investigate lipid diffusion and flows in our system. Fluorescence Recovery After Photobleaching (FRAP) assays revealed no significant difference in the recovery times across different cell regions at the ventral membrane (Figure S5A, note that FRAP t_1/2_ does not change on the adhesive and non-adhesive areas of the edges). We further tested the presence of flows using FRAP, bleaching a line tangential to the cell edge and tracking if the fluorescence minima moved during recovery. These assays showed that intensity minima moved towards the cell edge (Figure S5B). However, this apparent lipid flow did not change in the absence of actin dynamics (Figure S4F-G, S5B, JLY treatment). FRAP data can be interpreted as resulting from either a ventral plasma membrane flow directed towards the cell edges that is non-dependent on actin dynamics, or from the existence of a diffusion barrier at the leading edge. While the presence of a diffusion barrier at the leading edge was proposed by previous studies,^80^ we aimed at discriminating between flows and diffusion barriers with other means than bleaching.

For this, we employed fluorescence fluctuation spectroscopy (FFS) and spatiotemporally investigated lipid dynamics without photobleaching by concomitant autocorrelation function (ACF) and pair correlation function (pCF) analysis across line scan FFS measurements.^81–83^ In particular, single-channel line scans across the ventral membrane of patterned cells stained with the plasma membrane dye CellMask were acquired at a rate faster than the rate at which the membrane dye molecules diffuse (Figure S5C-D). Then the local versus long range mobility of the diffusing dye molecules across the membrane was extracted by an ACF and pCF analysis of the acquired fluorescence fluctuations with themselves versus at specific distances in the forward or reverse direction (Figure S5E, see Methods). We positioned the line scans crossing the pattern boundaries, thus across regions with a sharp decrease of the Flipper-TR fluorescence lifetime (Figure S5G). The ACF analysis showed that lipid mobility does not change between the adhesive and non-adhesive areas (Figure S5G-H). The diffusion values are consistent with point FCS measurements of the same lipid probe from the literature (⟨D_cross control_⟩ = 2.2±2.7μm^2^/s; ⟨D_ring control_⟩ = 1.9±2.6μm^2^/s)^84^. Under JLY treatment, we noticed a weak trend decreasing overall diffusion and transport (⟨D_cross JLY_⟩ = 1.7±2.0μm^2^/s; ⟨D_ring JLY_⟩ = 1.5±2.0μm^2^/s). The pCF analysis revealed an absence of membrane lipid flow (Figure S5G, no significant differences between forward and retrograde transport). From these data, we concluded that patterned cells do not display a steady-state lipid flow at their ventral membrane, despite the presence of membrane tension gradients. This is consistent with the fact that membrane tension gradients are maintained through time, as they do not dissipate by lipid flows.

### Substrate rigidity and adhesiveness modulate membrane tension

Our results in patterned cells indicate that tension gradients can exist in non-migrating cells due to adhesion patterns and actin polymerization. Therefore, this proves that tension gradients are not exclusive to migrating cells. We next wondered if they were exclusive to adherent cells. To address this, we examined whether membrane tension gradients are present in non-adherent migrating cells which usually migrate through bleb expansion. We confined HeLa cells to trigger leader bleb migration—a mode of migration independent of specific adhesion, but where a gradient of cortical actin orients blebbing at the leading edge of the cell.^85,86^ Cells were confined for over an hour to ensure the predominance of the leader bleb phenotype (Figure 5A).^87^ Our findings indicated that Flipper-TR lifetime gradients are absent at the protruding front of leader bleb migrating cells (Figures 5B and 5C). The lifetime was only marginally increased at the rear of the leader bleb, adjacent to the uropod (i.e. the adhesive rear of the cell training behind the leader bleb). Nevertheless, the magnitude of these lifetime changes was negligible compared to the gradients observed in adherent cells. This shows that adhesion is necessary to create a tension gradient, suggesting that tension gradients are not essential for migration.

**Figure 5:**
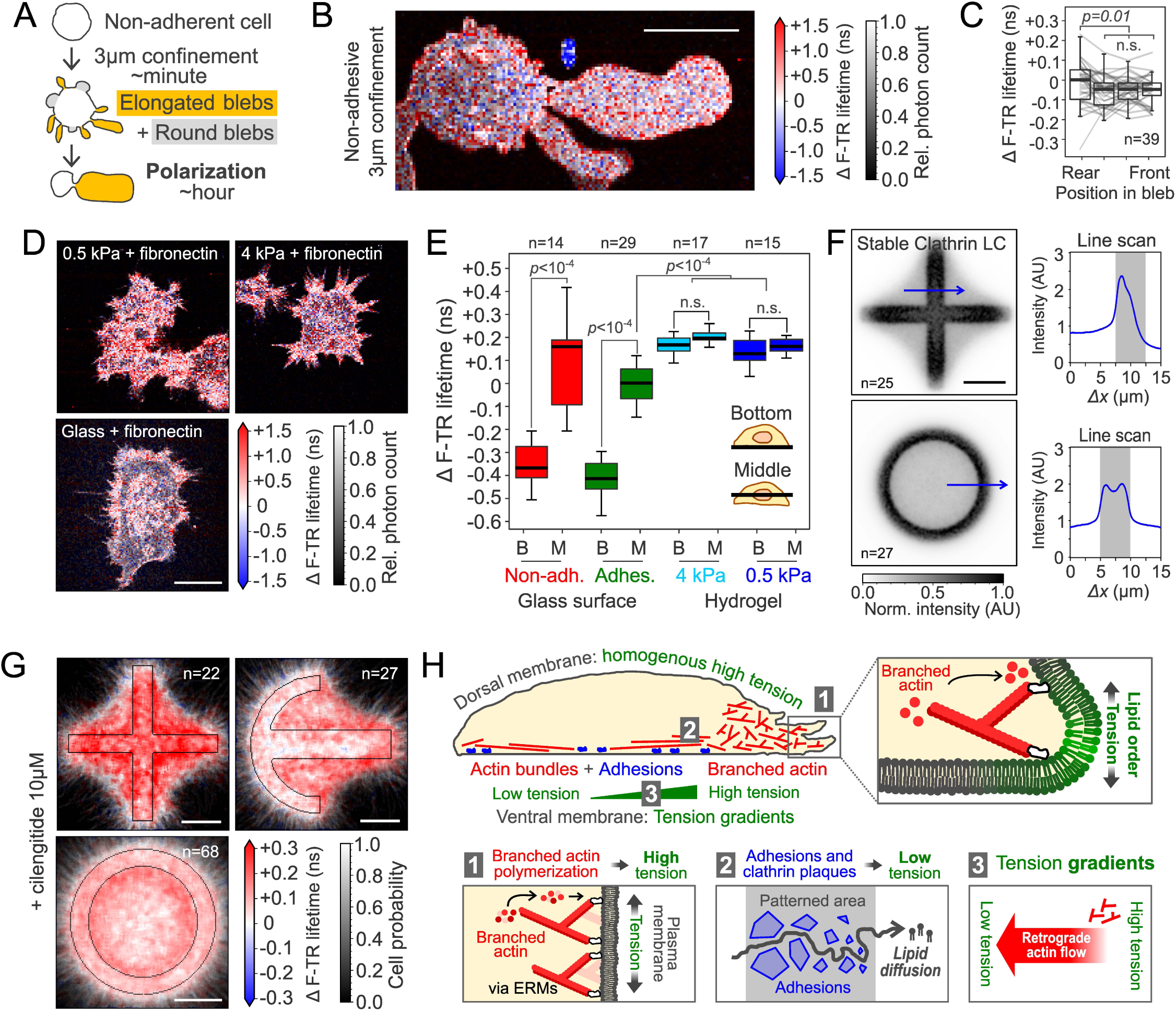
Substrate rigidity and adhesiveness modulate membrane tension. **A**: Diagram of confinement experiment showing cell morphology and bleb formation before polarization into a leader bleb. **B**: Representative FLIM image of 3μm-confined HeLa cells on pll-g-PEG (non-adhesive) stained with Flipper-TR. Lifetime is shown as Δτ from the middle plane average of cells on glass. **C**: Median, quartile distribution, and individual traces of Flipper-TR lifetime in stable blebs at different sections. Welch’s p: 0.013 or n.s. **D**: Representative FLIM image of HeLa cells on different substrates stained with Flipper-TR. Lifetime shown as Δτ from the middle plane average of cells on glass. **E**: Median and quartile distribution of Flipper-TR lifetime in HeLa cells on non-adhesive confinement, fibronectin-coated glass, and fibronectin-coated hydrogels (4.0 and 0.5 kPa). Welch’s p<10^-3^ or n.s. **F**: Average clathrin light chain intensity and line scans (blue lines) for cross (top, n=25) and ring (bottom, n=27) patterned cells. Patterned area in grey. **G**: Difference in Flipper-TR lifetime of cilengitide-treated cells from control average for cross, crossbow, and ring-patterned cells. **H**: Conceptual model of membrane tension gradient maintenance. The diagram shows tension variation across the cell, highlighting high tension on the dorsal side and gradients on the ventral side. Key factors include branched actin filaments (red), cellular adhesions, and clathrin plaques (blue) acting as barriers, creating tension gradients and directing lipid flow, crucial for cellular processes like movement and signaling. **B, D, F, G**: Scale bar, 10 µm. A: 1 min.

Since adhesion is essential to create tension gradients in the plasma membrane, we hypothesized that the modulation of adhesion strength could change those gradients. Specifically, we wondered if adhesion could be responsible for the differences in lifetime observed between the bottom and middle planes of the cells (Figures S3B-C and S4E). This difference was persistent despite various pharmacological treatments perturbing the actin cytoskeleton, indicating that it was not driven by actin dynamics. Considering leader bleb cells exhibit the same shift between bottom and middle plane lifetimes as cells adhering to fibronectin-coated glass, we assume the ventral-to-dorsal shift in average lifetime does not depend on specific molecular interactions. As substrate stiffness is known to strongly affect the adhesion strength of cells,^88^ we plated HeLa cells on substrates of different stiffnesses (Figure 5D) and measured the average Flipper-TR lifetime at the ventral and dorsal membranes, i.e., bottom and middle planes (Figure 5E). Interestingly, cells plated on fibronectin-coated soft polyacrylamide (PAA) hydrogels (E = 0.5 and 4.0 kPa, Methods), did not exhibit any differences in the average Flipper-TR lifetime between middle and bottom planes (Figure 5E). This was due to the lifetime at the bottom membrane of cells increasing on soft substrates with respect to the bottom membranes of cells plated on glass. Moreover, we observed an attenuation of lateral lifetime gradients on soft hydrogels. These data confirmed not only that adhesion per se is essential for the generation of tension gradients, but also that adhesion strength, which scales with substrate stiffness,^88^ is also critical.

To further test the role of adhesion strength in the generation and sustaining of plasma membrane tension gradients, we next aimed at pharmacologically reducing adhesion. Because tension gradients did not correlate with focal adhesions, and because removing focal adhesions may lead to complete detachment of cells, we investigated other adhesive structures. Clathrin plaques, distinct from transient clathrin-coated pits, are adhesive structures that rely on αvβ5 integrin-ECM interactions and are present in some cell lines when bound to high-rigidity substrates.^89^ We patterned HeLa cells transiently expressing clathrin light chain fused with GFP and quantified the localization of the stable clathrin population using TIRF microscopy. Clathrin plaques colocalized with low-lifetime zones, appearing at the inner edge and at the edges of the cross pattern (Figure 5F). Treatment with cilengitide, which inhibits αvβ5-integrin-mediated adhesions, resulted in an increase of Flipper-TR lifetime at the areas surrounding the patterns (Figure 5G), like the effect of plating cells on low-rigidity substrates. Thus, reducing adhesion strength by releasing molecular bounds flattens membrane tension gradients.

## Discussion

By using Flipper-TR as a mechano-responsive molecular probe, here we established how branched actin networks and cellular adhesions play a critical role in generating and sustaining spatial tension gradients within the ventral plasma membrane of adherent cells. Also, employing *in vitro* experiments with model membranes, we could establish the exact nature of the coupling between Flipper-TR lifetime and membrane tension. Therefore, this work validates the use of Flipper-TR as an indicator of membrane tension and shows the influence of lipid composition on the relation between tension and lipid packing. In particular, we describe how branched actin networks and membrane-to-cortex attachment are essential for tension buildup and tension propagation, respectively.

The differences between the upper and lower values of Flipper-TR lifetime gradients are in the range of 0.20 ns. From the calibration between flipper-TR lifetime and effective tension in HeLa,^24,90^ we can evaluate *Δσ_eff_* to be in the range of 0.25 mN.m^−1^. Therefore, we estimate the tension to drop to be about 30-40% from the resting *σ_eff_* of 0.1-0.6 mN.m^−1^ found in all cell types studied, which is far from being a negligible tension gradient. Within the same range, an increase of 30% higher at the leading edge compared to the trailing edge was previously measured in fish keratocytes by other means.^12^

Some studies proposed that membrane tension propagates very fast, therefore homogenizing tension values,^32^ while others have proposed that tension gradients can be maintained in polarized cells.^12,91^ Our work reconciles both ideas by proposing that tension propagation depends on the degree of membrane-substrate interaction. Therefore, contrary to what has been suggested,^35^ the existence of tension gradients does not depend on the cell type, but on the migration mode.^87^ At the ventral membrane, attached to the coverslip, we find strong inhomogeneities, which disappear at the apical membrane not attached to the substrate (Figure 5G, top panel). This allows us to propose a mechanism by which membrane tension gradients can be sustained in adherent cells (Figure 5G). Dynamic extension of the membrane by actomyosin turnover at the cell edge locally increases the tension. The propagation of tension towards the cell center is hindered by adhesion to the substrate and between the membrane and the actin cortex. The constant pulling force at protrusion sites combined with the decrease of tension at adhesions set the shape of the tension gradient. In the following, we discuss the consistency of this model with previous literature.

We propose that cells must adhere to generate and sustain tension gradients. A remaining question is which are the molecular principles at play to limit lipid diffusion. While rapid lipid diffusion can relax tension gradients, tension could be relaxed by other mechanisms, such as membrane unfolding or osmotic pressure release.^21,90^ Therefore, a reduction of lipid diffusion is not sufficient to create tension gradients, and assuming that hindered propagation of membrane flows is a signature of tension gradient may be fallacious.^34^ Also, lipid diffusion depends on which scale it is measured. At the nanometer scale, no major difference in mobility is observed in lipid diffusion^92^ between the ventral and dorsal membrane. At the submicron scale, lipid diffusion may be constrained by ‘corrals’ created by the mesh of cortical actin,^93^ by transient actin asters driven by Arp2/3^94,95^, or by lipid nanodomains. The two last modes are very dynamic, so their contribution to the long-range flow barrier is less obvious. But it is really at the cell scale that lipid flows may counteract or create membrane tension gradients. This may explain discrepancies found in the literature on how lipid flows and membrane tension gradients are coupled, as the mobility of lipids is usually investigated at a molecular scale, while gradients are always evidenced at the cellular scale.

Regarding the role of actin, our data shows that branched actin both increases membrane tension at the leading edge and plays a role in its propagation. There is extensive evidence linking branched actin networks to membrane tension increase, often via ERM proteins.^13,21,32,35^ What molecular mechanism mediates branched actin tension increase? On one hand, the branched actin nucleator Arp2/3 must bind to membrane-bound WAVE to add new filaments,^96^ linking actin polymerization and membrane binding. However, the specific properties of branched actin networks that promote tension increase might instead originate from their architecture, which confers them the ability to adapt density and pushing angle to the load to maintain polymerization.^17,97,98^

Overall, our study demonstrates that membrane tension gradients in adherent cells arise from the interplay between actin dynamics, lipid diffusion/flow constraints, and cell-to-substrate adhesion. By visualizing tension gradients, our work links the physical properties of the plasma membrane with cytoskeletal dynamics and understands how cell membrane tension gradients are formed, opening new avenues for investigating their role in cellular processes such as receptor signaling, phagocytosis, cell polarity or cell differentiation. For example, we speculate that membrane partitioning and self-organized criticality may allow lipid organization to respond to small perturbations like tension changes,^46^ impacting allosteric interactions at the membrane and cell polarity signaling.

### Limitations of the study

The limitations of our study stem from the use of fluorescence lifetime imaging microscopy (FLIM) on a scanning confocal setup. This technique is slow compared to the speed of actin dynamics, and especially to membrane diffusion (the acquisition time of single cells was ≈ 30s), which means that images of single cells are in fact averaged over times during which both actin and lipids move. Other FLIM techniques such as gated and modulated CCD image sensors might enable one to go faster and obtain images with higher time resolution.^99^ While our results from MALDI-MSI on micropattern cells agree with lipid distributions expected from membrane tension gradients *in vitro*, it is important to note that MALDI-MSI requires fixation and quantifies lipids from all membranes, not just plasma membrane, factors that may interfere with the experimental interpretation. These factors may limit the use of MALDI-MSI to study lipid compositional gradients in live cells.

## Acknowledgments

We thank Javier Espadas for training on supported lipid bilayers and useful feedback, Martin Lenz for discussions on membrane flows, Julie Miesch and Anne-Laure Boinet for TIRF microscope training, and Vincent Mercier and Chloé Roffay for FLIM confocal microscope training. Thanks to Rafael Caetano, Frédéric Humbert, and Roux lab members for technical assistance and feedback. Special thanks to Mireia Andreu Calvo, Guillaume Pernollet, Vincent Mercier, and the ACCESS platform for micropatterning training. We thank Henry De Belly, Matthieu Piel, Mathieu Dedenon, Chloé Roffay and Larisa Venkova for critical reading. A.R. acknowledges funding from the Swiss National Fund for Research grant numbers #CRSII5_189996 and #310030_200793 and the European Research Council Synergy grant number #951324-R2-TENSION. A.M. acknowledges funding from the Human Frontiers of Science Program (grant number LT000762/2020-L). P.G. acknowledges support from the Human Frontiers of Science Program (grant number LT-000793/2018-C). C.A. acknowledges funding from the DIP of the Canton of Geneva, SNSF (31003A_182473 and TMSGI3_211433).

## Author contributions

JMGA: Conceptualization, Methodology, Software, Formal Analysis, Investigation, Data Curation, Writing – Original Draft, Writing – Review & Editing, Visualization, Project Administration. AR: Conceptualization, Methodology, Writing – Original Draft, Writing – Review & Editing, Supervision, Project Administration, Funding Acquisition. AM: Software, Formal Analysis, Visualization. EH: Software, Formal Analysis, Supervision, Funding Acquisition. JS: Formal Analysis. PG: Investigation, Writing – Review & Editing. CT: Investigation, Writing – Review & Editing. AC: Investigation. GD: Supervision, Funding Acquisition. CA: Supervision, Project Administration, Funding Acquisition.

## Declaration of Generative AI and AI-assisted technologies in the writing process

During the preparation of this work, the authors used Grammarly and ChatGPT in order to improve readability in minor parts of the text. After using this tool, the authors reviewed and edited the content and take full responsibility for the content of the publication.

## STAR Methods text

### RESOURCE AVAILABILITY

#### Lead contact

Further information and requests for resources and reagents should be directed to and will be fulfilled by the lead contact, Aurélien Roux.

#### Materials availability

This study did not generate new unique reagents.

#### Data and code availability

All original microscopy images and datasets shown in the figures, and all original code and associated datasets have been deposited at an open repository and will be made publicly available as of the date of publication. Any additional data related to this paper will be shared by the lead contact upon request.

Figshare data DOI: https://doi.org/10.6084/m9.figshare.26304718

Any additional information required to reanalyze the data reported in this paper is available from the lead contact upon request.

### EXPERIMENTAL MODEL AND STUDY PARTICIPANT DETAILS

#### Mammalian cell lines

Human cervical adenocarcinoma cells HeLa-Kyoto stably expressing myosin IIA (Myh9)-GFP and LifeAct-mCherry or Myh9-GFP, or LifeAct-mCherry and the plasma membrane-targeting CAAX box fused to GFP, or TALEN-edited ActB fused with GFP (Cellectis, Paris, France), or HeLa, Cos7 and U2OS cells with no stable marker were cultured in DMEM GlutaMAX medium (Gibco, #61965-026) supplemented with 10% FBS (Gibco, #P30-193306A) and Penicillin Streptomycin (Thermofisher, #15140-122) at 37°C and 5% CO2. RPE1 cells were cultured in DMEM / F-12 (Gibco) supplemented with 10% FBS and 1% Pen-Strep at 37°C in a 5% CO_2_ incubator.

All cell lines were tested for mycoplasma contamination using Mycoplasmacheck PCR Detection (Eurofins, #50400400), and treated once a year with Plasmocin profilactic (InvivoGen, #ant-mpp) and treatment Mycoplasma Elimination Reagents (InvivoGen, #ant-mpt).

#### Zebrafish keratocytes

Keratocyte cells were extracted from scales of zebrafish (*Danio Rerio*) while anesthetized with Ethyl 3-aminobenzoate methanesulfonate (Tricaine, SIGMA, E10521). 3-5 scales were extracted from each fish and placed on a glass-bottom dish containing Leibovitz’s medium (L-15, ThermoFisher #21083027), supplemented with 10% fetal bovine serum and 1% penicillin/streptomycin antibiotic solution. The scales were subsequently sandwiched between the dish’ glass and a coverslip and placed in an incubator at 28°C. Keratocytes were allowed to spread out from the scales for 12h. Subsequently, cells were washed twice with PBS and detached from the plates by treating them with trypsin-EDTA for 2-5 minutes. In order to remove the excess trypsin, cells were centrifuged in culture medium at 1000 rpm for 3 minutes. After discarding the supernatant, cells were resuspended in medium and plated in glass-bottom dishes for imaging. This protocol was carried out by trained personnel at the Zebrafish Core Facility of the medical center of the University of Geneva.

### METHOD DETAILS

#### DNA constructs and transfections

Cells were transfected with plasmid DNA using Lipofectamine 3000 reagent (Thermofisher #L3000001) transiently, according to manufacturer’s protocol, and imaged from 24 to 72 hours after transfection. The following expression vectors were used for plasmid DNA transfections: VAMP7-pHLuorin-pcDNA3 (gift from Thierry Galli, Inserm, France), eGFP-Clathrin Light Chain C1 (gift from P. De Camilli, Yale University, USA).

#### Drug treatments and chemical probes

The following pharmacological inhibitors and chemical compounds were used: 10 µM of ezrin inhibitor NSC668394 (Sigma #341216), 20 µM of Rho Kinase inhibitor Y-27632 (Sigma #Y0503), 8 µM of actin filament stabilizer Jasplakinolide (Cayman, #CAY-11705-100), 5 µM of actin polymerization inhibitor Latrunculin B (Sigma, #428020), 1 µM of actin polymerization inhibitor Latrunculin A (Sigma, #L5163), 85 µM of Arp2/3 actin polymerization inhibitor CK-666 (Sigma #SML0006), 10 µM of α_V_ integrins inhibitor Cilengitide (Sigma #SML1594), and 10 μg/ml actin inhibitor Cytochalasin B for cell enucleation experiments (Sigma, #250255). Cells were imaged 30 minutes after treatment. CellMask Green plasma membrane stain (Sigma, #C37608) and Flipper-TR live cell fluorescent membrane tension probe (Lubio Science and Spirochrome, #SC020) were used according to manufacturer’s specifications at 1:1000 from a stock dilution in DMSO, to a final concentration of 50 nM. SiR-actin staining (Spirochrome #CY-SC001) was performed at 1 µM during 30 minutes with 10 µM verapamil following the manufacturer’s instructions.

#### Cell fixation and staining

Adherent cells were fixed for 15 minutes with 4% paraformaldehyde in PBS. Cells were then permeabilized and blocked with the blocking buffer [5% Bovine Serum Albumin and 0.1% Saponin in PBS] for 30 min. Cells were washed three times with PBS before a 1 h incubation with Phalloidin-AlexaFluorPlus647 at 1:400 from a DMSO stock according to manufacturer’s instructions (Thermofisher #A30107), Anti-Vinculin Vinculin Monoclonal Antibody (7F9)-Alexa Fluor488 at 1:25 dillution (eBioscience Thermofisher #53-9777-82) on 1% Gelatin in PBS. Cells were then washed three times with PBS, labeled by NucBlue Fixed Cell Stain (DAPI, Thermofisher #R37606) for 10 min, and washed again three times with PBS. Once stained, the cells were imaged directly on the microscope via the saved localization.

For lipid localization, we stained cells with fluorescently-labelled bacterial toxins that recognize different sphingolipid headgroups: Shiga toxin 1a-Cy3 (ShTxB1a, at 1:50) binds to Gb3^100^, Cholera toxin-AlexaFluor488 (ChTxB, at 1:800) binds the ganglioside GM1^101^, Equinatoxin II-AlexaFluor647 (EqTx, at 1:200) binds sphingomyelins^102^ (all gifts from Giovanni D’Angelo, EPFL, Switzerland), and Filipin III at 25 μg/ml (Sigma, #F4767). The cells were fixed with 4% PFA, blocked in PBS containing 5% bovine serum albumin (BSA) without detergent, and incubated with fluorescently labelled B-subunit toxins for 1h.

#### Cell enucleation

The day before enucleation, U2OS and RPE1 cells detached with trypsin/EDTA (0.05% for 2 min, Gibco), and spread on a plastic polystyrene slide covered with bovine Fibronectin (FN, 10μg/ml, Sigma-Aldrich #F4759) in their respective culture medium.

During enucleation, plastic slides seeded with cells were incubated at 37°C for 15 minutes in the presence of 10 μg/ml Cytochalasin B (Sigma, #250255), then centrifuged at 15000 g for 40 minutes at 37°C in the presence of Cytochalasin. After centrifugation, to recover from the effects of Cytochalasin, the cells on plastic slides were washed with PBS, and transferred back to their respective drug-free culture media for 4 h at 37°C before use.

#### Migration time lapses of different mammalian cell lines

Mammalian cells were detached with trypsin/EDTA (0.05% for 2 min), then spread at a density not exceeding 10% confluence into 8-well chambered cover glass (Lab-Tek II) coated with bovine FN (10μg/ml) in their respective culture medium, 1 to 3 hours before the experiments.

Time-Lapse Phase-Contrast Microscopy Multisite microscopy of cells in 8-well chambered cover glass was performed in a humidified CO_2_ chamber with an Axio Observer Inverted TIRF microscope (Zeiss, 3i) and a Prime BSI (Photometrics) using a 10X objective (Zeiss, 10X). SlideBook 6 X64 software (version 6.0.17) was used for image acquisition. Cells were imaged using phase-contrast imaging, with a time-lapse of 5 minutes between frames for over 10 hours.

#### Confocal FLIM live imaging of Flipper-TR

During live FLIM acquisition, a corresponding phenol-red-free alternative of each culture medium was used. U2OS, HeLa and Cos7 cells were imaged in FluoroBrite DMEM (Gibco) during migration live experiments or Leibovitz’s L-15 (Thermo Fisher Scientific, #21083027) during micropatterning experiments in incubation at 37°C but without CO_2_. Media was supplemented with 10 % FBS and 1 % Pen-Strep while RPE1 cells were imaged DMEM/F-12 without phenol red (Gibco) supplemented with 10 % FBS and 1 % Pen-Strep.

Generally, cells were labelled for at least 10 minutes with FBS-free 50 medium containing 50 nM Flipper-TR. For supported lipid bilayers, the medium consisted on an aqueous buffer of 10mM HEPES at pH 7.4. After labeling with Flipper-TR or Flipper-TR and Hoechst, cells were imaged directly in glass-bottom microwell dishes at 37°C and 5% CO2. For migration assays, untreated and enucleated cells were detached with trypsin/EDTA (0.05% for 2 min), then spread at a density not exceeding 20% confluence into 35 mm glass-bottom microwell dishes (MatTek) coated with bovine FN (10μg/ml) in their respective culture medium, 1 to 3 hours before the experiments. After spreading, the untreated cells were labeled with Flipper-TR (Spirochrome) in their respective acquisition medium and incubated at 37°C for 10 min without probe washing. The Flipper-TR stock solution was composed by dilution of a 1 mM Flipper-TR in DMSO as previously described ^24^. After spreading, the enucleated cells were labeled by 1:1000 dilution of a stock solution of Flipper-TR and labeled by NucBlue Fixed Cell Stain (DAPI, Thermofisher #R37606) in their respective acquisition medium and incubated at 37°C for 10 min without probe washing.

The microscope was an inverted motorized microscope Eclipse Ti2 (Nikon) with a point scanning A1 confocal system from Nikon, equipped with a Plan Apo λ 100x oil objective with a NA 1.45 (Nikon, # MRD01905), a perfect focus system, a stage-top incubation chamber (Okolab) allowing long acquisitions. The imaging on hydrogels and comparing experiments was done instead with a water immersion 40x objective NA 1.15 (Nikon, # MRD77410) to allow for a higher working distance. The A1 confocal system was equipped with 4 excitation lasers at 405, 488, 561 and 640 nm. For fluorescence lifetime imaging (FLIM), we used a time-correlated photon counting system from PicoQuant integrated on Nikon Elements software. Excitation was performed using a pulsed 485 nm laser (PicoQuant #LDH-D-C-485) mounted on a laser-coupling unit, and a computer-controlled multichannel picosecond diode laser driver (PicoQuant, Sepia PDF 828) operating at 20 MHz. The emission signal was collected through a 600/50 nm bandpass filter using a gated PMA hybrid 40 detector and a multichannel time-correlated single photon counting system (PicoQuant, MultiHarp 150).

Single frames resulted from the integration of 2-10 frames depending on the sample and the confocal scanning area, to a total count between 10 and 50 photons per pixel. Frame rate during time lapse imaging in mammalian cells had to be set at 10 minutes to avoid phototoxicity and arresting actin dynamics.

#### Confocal live imaging for fluctuation correlation spectroscopy

Regarding fluctuation correlation spectroscopy analysis, the microscope setup was the same as for FLIM. For both line scanning used for autocorrelation function (ACF) and paired-correlation function (pCF) analysis, pixel dwell time was set at 5.2 microseconds, line time at 960 microseconds, pixel size was set at 160 nm, giving a line length of 10 micrometers. We positioned the 10-micrometer confocal line scans crossing the pattern boundaries, thus across regions with a sharp decrease of the Flipper-TR fluorescence lifetime. For the ring pattern, the line scan sits at the cell edge in the region where retrograde flows are dissipating. The adhesive area is located on the outer side of the line scan. For the cross pattern, the line scan is located closer to the center of the cell, and the adhesive pattern is on the inner side of the line.

#### Preparation of supported lipid bilayers

For supported lipid bilayer experiments, we used: 18:1 (Δ9-Cis) PC (DOPC) (Avanti #850375), 18:1 PS (DOPS) (Avanti #840035), porcine brain SM (Avanti #860062), plant cholesterol (Avanti #700100) at the specified mol% ratios. Some lipid mixtures included 0.02 mol% Atto 647 1,2-Dioleoyl-sn-glycero-3-phosphoethanolamine (DOPE) (Sigma #67335). Phospholipids or cholesterol stocks were dissolved in chloroform at concentrations ranging from 1-10 mg/ml and stored at −80°C in Argon atmosphere. Lipid mixes were calculated for a final mass of 0.25 mg. The chloroform in the vials was evaporated under an Argon flow in a fume hood and lipid films were formed. The vials were then stored in a vacuum oven at room temperature overnight to remove any residual chloroform.

To prepare liposomes, the vials were brought to room temperature and rehydrated by adding aqueous buffer HEPES 10mM at pH 7.4 to a final concentration of 0.5 mg/ml of lipids and mixing. The liposome mixtures were placed on a parafilm surface. 50-micron silica particles (Sigma #904384-2G) were first diluted 3x with distilled water, washed three times, and deposed on the lipid droplets on parafilm. The liposome-beads aqueous mixtures were then stored in a vacuum oven at room temperature overnight to remove any residual chloroform. To prepare the spreading bilayer assay, culture-grade 35-mm glass-bottom dishes (Mattek #P35G-1.5-14-C) were plasma cleaned (Harrick Plasma #PDC-32G) for 2 minutes at high power. Immediately after, dried liposome-bead mixtures were scrapped from the parafilm surface, deposed on the plasma-cleaned surface, and rehydrated with 1ml of 10 mM HEPES pH7.4 buffer containing 50 nM Flipper-TR (Lubio Science and Spirochrome, #SC020)

#### TIRF-FRAP experiments on patterned cells

All total internal reflection fluorescence microscopy (TIRF) movies were recorded on an Olympus IX83 widefield microscope equipped with a 150×/NA1.45 objective and an ImageEM X2 EM-CCD camera (Hamamatsu) under the control of the VisiView software (Visitron Systems). The 488 nm and 561 nm laser lines were used for illumination of GFP- and mCherry-tagged proteins. Excitation and emission were filtered using a TRF89902 405/488/561/647 nm quad-band filter set (Chroma). Laser angles were controlled by iLas2 (Roper Scientific).

Fluorescence recovery after photobleaching (FRAP) experiments were performed using a custom-built set-up that focuses a 488-nm laser beam at the sample plane, on the Olympus IX81 widefield microscope described above. The diameter of the bleach spot or width of the bleach line was approximately 0.5 μm.

#### Chemical passivation and micropatterning

Glass surfaces passivated with Polyethyleneglycol (PEG) were prepared for protein micropatterning. First, glass bottom dishes (Mattek) were activated by exposing them to air plasma (Harrick Plasma, PDC-32G) for 3LJminutes. Subsequently, the dishes were treated with a 0.1 mg/ml poly-lysine (PLL) (Sigma) solution for 30 min and washed with 10mM HEPES buffer (pHLJ8.4). By using this same buffer, a solution of 50 mg/ml polyethylene glycol (PEG) (molecular weight 5,000) linked to a succinimidyl valerate group (SVA, Laysan Bio) was prepared and applied to passivate the PLL-coated surface for 1.5LJh. Finally, dishes were washed with PBS 3 times.

Micropatterns were generated by using the system PRIMO (Alvéole), mounted on an inverted microscope Nikon Eclipse Ti-2. In the presence of a photo-initiator compound (PLPP, Alvéole) and DMD-generated patterns of UV light (375 nm), PEG is degraded. After illumination (1200LJmJ/mm) through a 20x objective, PLL is exposed. After rinsing with PBS, fibronectin (Calbiochem) was incubated at 50 μg/ml at room temperature for 5LJmin to coat the PEG-free motifs with the cell-adhesive protein. The excess of fibronectin was washed out with PBS. PBS was finally replaced by medium, and a suspension of cells was added at densities of roughly 10^5^LJcells per cm^2^. Samples were kept in an incubator at 37 °C and 5% CO2. After 10–30 min, non-adhered cells were washed out.

#### MALDI-MSI sample preparation

Cells were directly seeded on micropatterns with cross shape in complete media. After aspiration of media, cells were washed twice with PBS, followed by fixation in 0.25% glutaraldehyde for 15 minutes. For MALDI-MSI analyses, 150 μL of 2,5-dihydroxybenzoic acid (DHB), (30 mg/mL in 50:50 acetonitrile/water/0.1% TFA), were deposited on the surface of the samples using the automatic SMALDIPrep (TransMIT GmbH, Giessen, Germany).

#### Cell culture on hydrogels

Commercially-available EasyCoat hydrogels (Cell Guidance Systems, # SV3510-EC-0.5-EA and # SV3510-EC-4-EA) were used to control substrate rigidity. The hydrogels were pre-coated with fibronectin to facilitate cell adhesion. Briefly, hydrogels of specified rigidity (0.5 kPa and 4.0 kPa) were equilibrated in PBS-fibronectin solution (1 µg/mL, Sigma #F1141) for 1 hour to ensure uniform coating. Excess fibronectin was washed off with PBS before seeding the cells onto the hydrogels. Cells were cultured on these substrates under standard conditions, allowing us to investigate the effects of substrate rigidity on membrane tension and cell behavior.

#### Cell confinement

Cell confinement was achieved using a PDMS micropillar array mounted on a coverslip mounted on a PDMS device (named ‘suction cup’) trigged by a vacuum pump and pressure controller, following established methods^85,103^. Briefly, a SU8 photolithography mold was prepared with micropillars. This mold is used to create a PDMS confinement chamber of 3 micrometers in height, plasma-bonded to a 12-mm glass coverslip. The microfabricated confiner coverslips were treated with plasma for 1 min, and incubated with 0.5 mg/mL pLL-*g*-PEG (SuSoS, PLL(20)-g[3.5]-PEG(2)) in 10 mM pH 7.4 HEPES buffer for 1h at room temperature. Cells were trypsinized and seeded in a glass-bottom dish coated with PLL-g-PEG and the PDMS device was placed on top, creating a confined environment between the glass and the PDMS coverslip. This setup allowed us to trigger leader-bleb migration by confining the cells to a 3-micrometer height, promoting a non-adhesive environment conducive to the desired migration behavior.

### QUANTIFICATION AND STATISTICAL ANALYSIS

#### Statistical analysis of experimental data

Statistical analyses for all experiments were performed in Python 3 (NumPy and SciPy libraries) or Microsoft Excel. Statistical data are presented as median or mean ± standard deviation or interquartile range. For each panel, sample size (*n*), experiment count (*N*) statistical tests used, and *P* values are specified in the figure panel and/or legends. Plots were made using the SciPy and matplotlib Python libraries.

#### Basic image processing

Basic image analysis and format conversion was performed on ImageJ/Fiji software. Images were imported from NIS Elements (.nd2 format) using the BioFormats plugin.

#### Migration parameters and cell morphology of mammalian cell lines

Individual cells were tracked semi-automatically by random selection of cells from video images and manual tracking of migration pathways using the Manual Tracking function in ImageJ. At least 50 cells were tracked for each cell type (n ≥ 50). The cell migration tracking data were analyzed as described previously^104^.

*Cell aspect ratio* was calculated from binary masks using Image/Fiji by fitting ellipses to the cell shape.

*Protrusion area* in control patterned cells was segmented manually from intensity images.

#### General considerations about the different Flipper-TR lifetime estimates

Flipper-TR is a mechano-responsive molecular probe that changes its fluorescence properties in response to changes in membrane packing, making it an invaluable tool for studying the mechanical properties of the plasma membrane. Upon insertion to the membrane, Flipper-TR’s fluorescence lifetime changes with the tension in the lipid bilayer. Generally, the fluorescence lifetime is the average time the molecule spends in the excited state before emitting a photon and returning to the ground state. To calculate the fluorescence lifetime of Flipper-TR, we used a time-correlated single photon counting (TCSPC) device as described above. This technique measures the time between the excitation pulse and the subsequent photon emission, producing as an output the histogram of photon arrival times. The distribution of these arrival times can be analyzed to determine the fluorescence lifetime.

Due to the photochemistry of the molecule, Flipper-TR photon arrival time distribution follows a bi-exponential decay *l*(*t*), in the form: 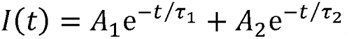, where *A_i_* and τ*_i_* represent the amplitude and the decay of a given exponential *i*, respectively. Previous studies have employed different methods to estimate the Flipper-TR lifetime. One common approach is to fit the fluorescence decay curve to a biexponential decay model, which accounts for the presence of two distinct exponential decay components ^24^. From this model, researchers can extract either the larger exponent (i.e.: τ_1_) or calculate an intensity-weighted average of the two exponents, 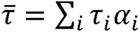 ^90^. These methods can provide detailed insights into the Flipper-TR lifetime, including a X^2^ used to judge the goodness, a greater precision, and the ability to decompose complex lifetime distributions, but they require more computational resources and can be more sensitive to noise and initial parameter estimates. Moreover, even at high photon counts, the ability to determine the precise values of *α_i_* and τ*_i_* by a biexponential fit can be hindered by parameter correlation ^105^. To avoid problems involved with fitting, other works have used phasor plots have been used to measure changes in Flipper-TR lifetime ^106,107^. Phasor plots offer a graphical representation of the fluorescence decay characteristics, simplifying the analysis of complex decay patterns. By mapping each pixel to specific G/S coordinates on the phasor plot, one can visually assess changes in membrane tension and identify different lifetime components without assuming any feature of the fluorescence decay, such as the number of exponents. This is particularly useful when using complex samples or Flipper variants where the photochemical properties have not been fully characterized.

For this work we have instead relied on the barycenter of the lifetime distribution (i.e.: mean photon arrival time or “fast lifetime” calculation) to estimate fluorescence lifetime. As the instrumentation induces a delay in the photon arrival, the average lifetime equals time span from the barycenter of the instrument response function (IRF) to the barycenter of the decay in a pixel-wise manner, as defined by our TCSPC manufacturer (PicoQuant). This method is simple, straightforward, and computationally efficient, especially reliable at low photon counts. The calculation yields a single lifetime exponent, that we refer to as “average Flipper-TR lifetime” τ̅, numerically equivalent to the intensity-weighted average of the two exponents coming from a fit. The need for a robust lifetime estimator at low photon counts was very important for this work as we were highly limited by sample illumination. Protrusion dynamics are particularly sensitive to phototoxicity and even relatively low illumination can arrest actin polymerization, so the photon counts were generally too low to be able to fit in a robust manner.

#### Flipper-TR lifetime estimation

##### Processing and plotting of FLIM images

FLIM images acquired on NIS software consisted of a channel containing average lifetime values and another with photon counts per pixel. The images were exported as .tif and processed on a custom-made Python script. Lifetime images were applied to a 2D median filter with a kernel size of 3 pixels (corresponding to 0.75 μm). In the exported lifetime images, the rainbow color scale in each pixel represents the values of the decay times. On top of the rainbow color scale, a dark mask is applied to represent photon counts (i.e.: black for no counts). This allows for a proper visual representation of the data because it masks the background where lifetime values are just due to noise.

##### Flipper-TR lifetime region-of-interest averages

When averaging pixels, the lifetimes of each pixel τ*_i_* were weighted by the photon count *n_i_*, so that 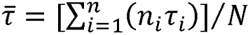. Different regions corresponded to specific criteria, explained as follows. In keratocytes, front/rear regions were manually segmented to cover approximately one-fourth of the cell each. In migrating mammalian cells, front/rear regions were defined as 60° angular sections extending 0-10 μm from the edge using a custom python script. Front and rear polarity angles were manually determined based on cell morphology, not lifetime values. For supported lipid bilayers, only the 100 μm closest to the advancing edge in the field of view was considered, using a custom python script. Front and rear regions were defined as the 20% closest and most distant areas from the advancing edge, respectively.

##### Flipper-TR lifetime distribution and ROIs in patterned cells

Generally, FLIM images were filtered and values under a threshold photon count of 20 photons were defined as NaN and not considered for analysis. For overall distribution, i.e. the lifetime spatial probability map, cells were aligned according to the patterned regions as previously described^108^, and a pixel-wise average was calculated using a custom python script. For cells adhered on micropatterns, different regions were defined to report spatial lifetime gradients: bottom/middle planes; cell body/protrusions; adhesive/non-adhesive; high/low. For the bottom plane there was no further segmentation, but for the middle plane (3 microns above the glass surface) a manual selection was used to segment the plasma membrane. Cell body and protrusions were manually segmented. Adhesive and non-adhesive regions were defined based on the fluorescence from micropatterns. Finally, to calculate the distance map 2D histogram, distances to the selected boundaries were defined per pixel using binary masks and the ‘distance map’ function in ImageJ/Fiji. Then, pixels were ordered and averaged to form a 2D histogram. To define the high and low regions we first defined the pixels were 99% of control cells were present. From this, we defined a ‘high’ region, containing the highest two lifetime deciles, and a ‘low’ region, containing the lowest 2 lifetime deciles of the control lifetime spatial distribution. These two regions of interest were applied to other conditions to represent lifetime gradients.

*Flipper-TR lifetime spatial decays* in mammalian cell lines were calculated using a custom python script based on distances to the cell edge and position in the micropattern. For radial/linear gradients, a 5 micron-wide region was defined and the average lifetime calculated per micron.

*Flipper-TR lifetime as a function of edge position* in migrating cells was calculated using a cell mask based on photon counts and, from a mask, a distance map to the edge. Then, pixels were ordered and averaged to form a 2D histogram.

##### Flipper-TR lifetime as a function of edge velocity

The edge velocity was calculated using a custom python script. For mammalian cells, the edge displacement was calculated by segmenting the cell contours at each timepoints and calculating the differences between the distance maps. For supported lipid bilayers, the images were manually cropped into rectangular regions. The velocity at each timepoint (μm/min) was defined as the net increase of area (μm^2^), divided by the frame rate (min) and the crop width (μm).

##### Flipper-TR lifetime in protruding/retracting cell regions

The condition to classify edges intro protruding/retracting was a velocity higher/lower than 0.2 μm/min. From this, angular sections extending 0-10 μm from the edge were defined.

##### Flipper-TR lifetime differences to control distributions

For some drug treatments, instead of the absolute Flipper-TR distributions, the difference to the control was instead plotted. First, the average control distribution was calculated on a pixel-wise basis. Then, these control values were subtracted to the drug-treated images on a cell-by-cell basis and the average differences plotted.

*Flipper-TR lifetime in hydrogels and non-adhesive migration* was instead calculated using the SymPhoTime 64 software (PicoQuant) to fit fluorescence decay data (from manual regions of interest) to a dual exponential model (n-exponential reconvolution) after deconvolution for the instrument response function calculated by the software. Lifetime was then expressed in difference in nanoseconds from the reference value, which is the lifetime at the middle plane of cells adhered to glass.

#### Particle image velocimetry of actin flows

##### PIV analysis

Movies were acquired at a 2-frame-per-second rate and segmented using Ilastik^68^ to define cell boundaries. Pixels outside cell boundaries were not considered for the PIV calculation. PIV vectors were calculated on a custom macro based on the ‘Iterative PIV’ ImageJ/Fiji plugin^109^, using approximately an XY grid of 1.25 μm and a search area of 1.5 μm. Importantly, even though we used the ‘Iterative PIV’ plugin, the PIV was calculated on a single window basis, so there is no a priori correlation between neighbor vectors.

#### Quantification of membrane events

*Stable clathrin coverage* was calculated from a minimum time projection of TIRF movies over 5 minutes at a 1 frame/second rate. This projection was then binarized to display the sites where the clathrin signal was stable over 5 minutes on a cell-by-cell basis.

*Dynamic clathrin coverage* was calculated in an analogous manner. First, we obtained a binary mask resulting from a maximum projection of the clathrin signal. Then we subtracted to this mask the stable clathrin sites. This yields a mask containing the sites where the clathrin signal was dynamic at some point over 5 minutes on a cell-by-cell basis, excluding the clathrin plaques where the signal was stable.

*VAMP7 exocytosis events* were manually selected from 5-minute TIRF movies of VAMP7 transfected and patterned cells.

Finally, the probability density distributions were plotted as a function of the distance to the center of the pattern and normalized to integrate to 1.

#### Actin-based segmentation of cell regions

To determine the effect of actin organization on Flipper-TR lifespan. Phalloidin-A647 images of each cell were segmented into three regions; Lamellipodium, Lamella, and Cell body regions. Binary segmentations were generated using the pixel classification process in Ilastik^68^. Segmentation masks were then applied to the Flipper-TR lifetime image acquired before fixation, to calculate the average Flipper-TR lifetime in each type of region for each cell.

#### Fluctuation correlation spectroscopy analysis

To study lipid diffusion in our system, we employed fluorescence fluctuation spectroscopy (FFS). This involved first construction of single-channel FFS line scans acquired across the ventral membrane of patterned cells stained with the plasma membrane dye CellMask into kymographs (x-dimension represents distance (i.e., 64 pixels), y-dimension represents time (i.e., 100,000 lines)) and application of a detrending algorithm to remove slow timescale artefacts (e.g., photobleaching). Then application of the autocorrelation function (ACF) on a pixel-by-pixel basis (*δr* = 0) to three spatially distinct sections across the kymograph (i.e., pixels 1-21, 22-42, and 43-63), which alongside fitting of the resulting ACFs to a 2-component diffusion model, enabled recovery of the average number of moving molecules (*G*_0_) as well as their local diffusion coefficient (*D*_0_) to be mapped across the non-adhesive versus adhesive coating and the interface in between (Figure 5E, right panel). And finally, application of the pair correlation function (pCF) on a pixel-by-pixel basis (*δr* = 5) across the entire kymograph (i.e., pixels 1-59 from left to right and pixels 5-64 from right to left), which alongside fitting of the resulting pCFs to a Gaussian function, enabled recovery of the molecules direction dependent arrival time (τ) as well as their transport efficiency (i.e., *G_τ_*/*G*_0_) (Figure 5E, left panel). Published codes are also available at https://github.com/ehinde/Pair-correlation-microscopy.git repository.

#### Analysis of TIRF-FRAP experiments on patterned cells

FRAP experiments performed on MyrPalm-GFP patterned HeLa cells were analyzed as previously described^111^. Mean fluorescence values were measured from regions of interest representing the background, the cell, and the membrane. A custom-written Python script was used to subtract background fluorescence, correct for photobleaching, and normalize the values between 0 (corrected fluorescence immediately after photobleaching) and 1 (mean corrected fluorescence of 5 s before photobleaching). The recovery curves of individual experiments were aligned to bleach time and averaged. The average was fitted to a single exponential equation from which the mobile fraction and recovery half-time were calculated.

To calculate the velocity of lipid flows on line-FRAP experiments, fluorescence intensity was integrated over 11 microns (100 pixels) parallel to the bleaching line, and the minima were tracked for 12 frames (0.5 seconds per frame) using a custom-made Python script.

#### MALDI-MSI data analysis

MSI experiments were performed using AP-SMALDI5 AF systems that couple a Q Exactive orbital trapping mass spectrometer (Thermo Fisher Scientific, Bremen, Germany) with an atmospheric-pressure scanning-microprobe MALDI imaging source (AP-SMALDI, TransMIT GmbH, Giessen, Germany). The MALDI laser focus was optimized manually using the source cameras aiming at a diameter of the focused beam of 7 μm. For each pixel, the spectrum was accumulated from 50 laser shots at 100 Hz. MS parameters in the Tune software (Thermo Fisher Scientific) were set to the spray voltage of 4 kV, S-Lens 100 eV, and capillary temperature of 250°C. The step size of the sample stage was set to 5 μm. Positive ion mode measurements were performed in full scan mode in the mass range *m/z* 400-1600 with a resolving power set to R = 240000 at *m/z* = 200. Mass spectra were internally calibrated using the lock mass feature of the instrument.

The output of the MALDI-MSI was processed using a custom-made Python script. The output consists of a multi-dimensional .tif with 389 channels representing the peak intensity at each *m/z* bin and XY spatial coordinates at 0.5 microns per pixel. First, an integration of all counts allowed to identify pattern positions with cells and to align patterns with one another. Then, the relative intensity of peaks from specific lipid species was calculated over the total intensity of only the known peaks (112/389). For lipid species present in more than one peak, the relative abundances were added together. This yields a lipid fraction per pixel, per cell, of each lipid species. Lipid fractions were also normalized to analyze their spatial heterogeneity. To do so, lipid fractions from a given lipid species from all cells, all pixels were pooled together, and their mean and standard deviation were calculated. Lipid fractions were then subtracted from the mean and divided by the standard deviation.

## Supplementary video titles and legends

**Supplementary movie 1**, corresponding to Fig. 1. Expanding supported lipid bilayers (SLBs) with different compositions.

**Supplementary movie 2**, corresponding to Fig. 2. Confocal FLIM time lapses of migrating U2OS cells stained with Flipper-TR.

**Supplementary movie 3**, corresponding to Fig. 5. Actin dynamics in patterned HeLa Actin-GFP cells, examples of lamellipodium and stress fiber dominated phenotypes.

**Supplementary movie 4**, corresponding to Fig. 5. Actin dynamics in patterned HeLa Actin-GFP cells in control (+DMSO) versus JLY-treated conditions.

---

**Figure S1:**
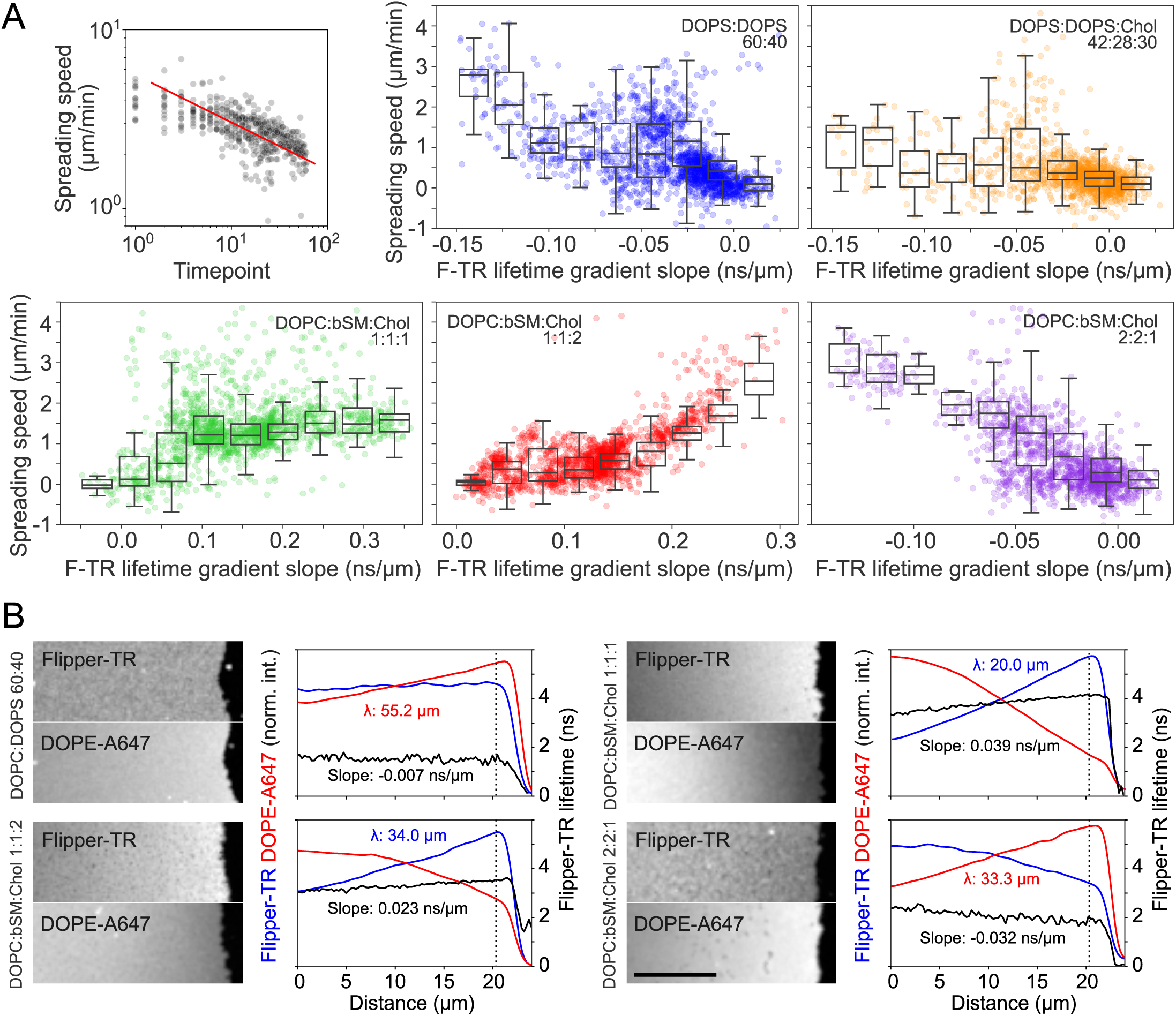
Flipper-TR reports membrane tension gradients in reconstituted membranes. **A**: Top left, bilayer spreading speed (µm/min) over time. Red line for visual reference. Rest, bilayer spreading speed (µm/min) as a function of linear fits of spatial Flipper-TR gradients (ns/µm) for different lipid compositions. Histograms of binned data overlaid in black. **B**: Left, fluorescence confocal images of Flipper-TR and DOPE-atto647 in spreading bilayers of different compositions. Right, spatial profile of Flipper-TR (blue) and DOPE-atto647 (red) fluorescence intensity, and Flipper-TR average fluorescence lifetime (black, ns). Exponential (for fluorescence intensity, λ) or linear fits (for lifetime, slope) overlaid.

**Figure S2:**
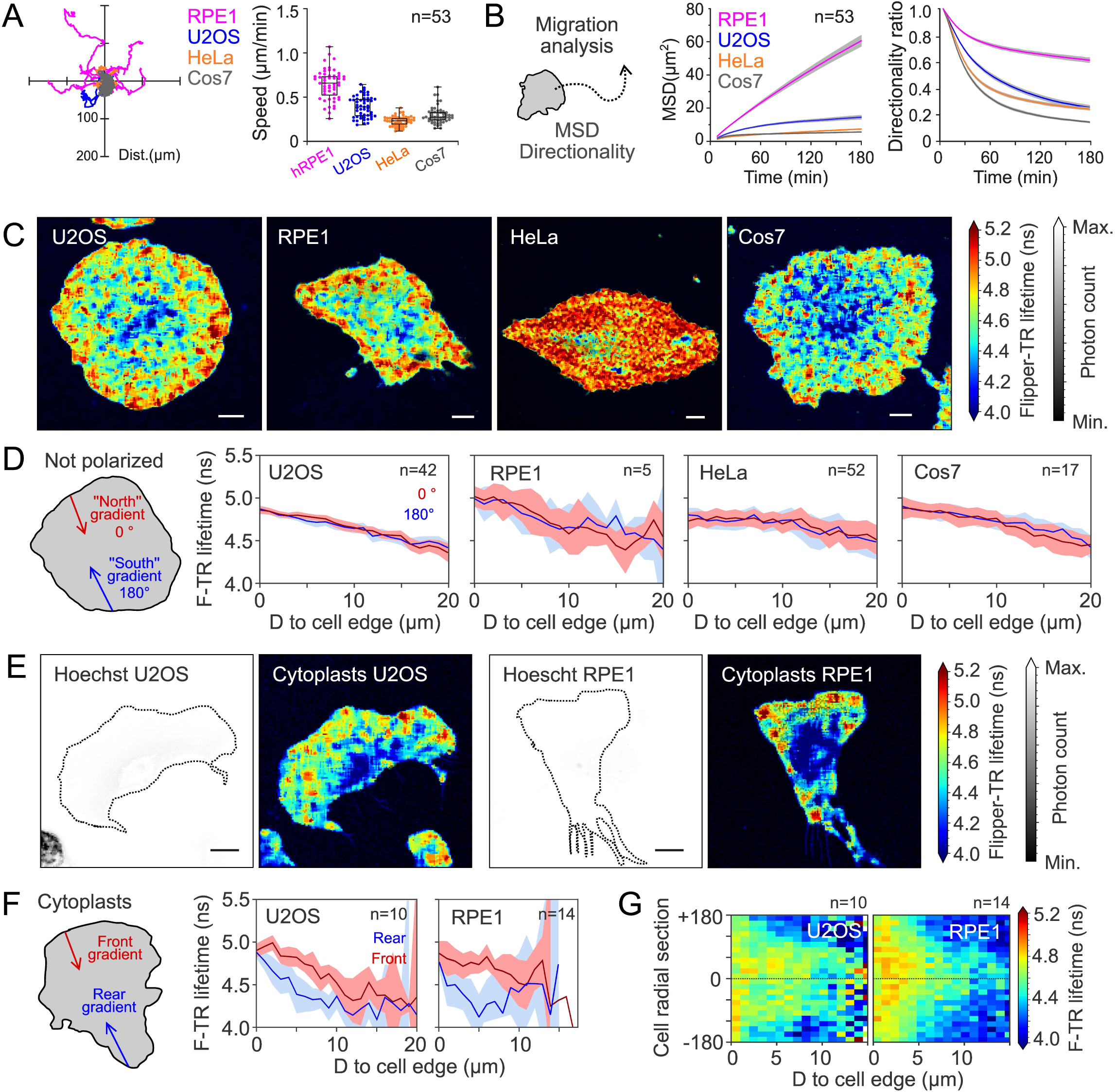
Leading-edge extension increases Flipper-TR lifetime in migrating cells. **A**: Mean square displacement (MSD, µm²) and directionality ratio as a function of elapsed time for RPE1 (magenta), U2OS (blue), HeLa (orange), and Cos7 (grey) cells, n=53. **B**: Instantaneous speed (µm/min) and trajectories over 180 minutes for RPE1 (magenta), U2OS (blue), HeLa (orange), and Cos7 (grey) cells, n=53. **C**: Representative image of non-polarized U2OS, RPE1, HeLa, and Cos7 cells labeled with Flipper-TR. **D**: Average Flipper-TR fluorescence lifetime as a function of distance from the edge at the front (red) and rear (blue) of non-polarized U2OS (n=90), RPE1 (n=37), HeLa (n=11), and Cos7 (n=15) cells. Line represents mean ± standard deviation. **E**: Representative image of U2OS and RPE1 cytoplasts labeled with Flipper-TR and Hoechst. **F**: Average Flipper-TR fluorescence lifetime as a function of distance from the edge at the front (red) and rear (blue) of U2OS (n=10) and RPE1 (n=14) cytoplasts. Line represents mean ± standard deviation. **G**: Average Flipper-TR fluorescence lifetime as a function of distance from the edge and radial position (front at 0°) of U2OS (n=10) and RPE1 (n=14) cytoplasts. Color indicates average Flipper-TR fluorescence lifetime (ns). **A,C,E**: Scale bar, 10 µm.

**Figure S3:**
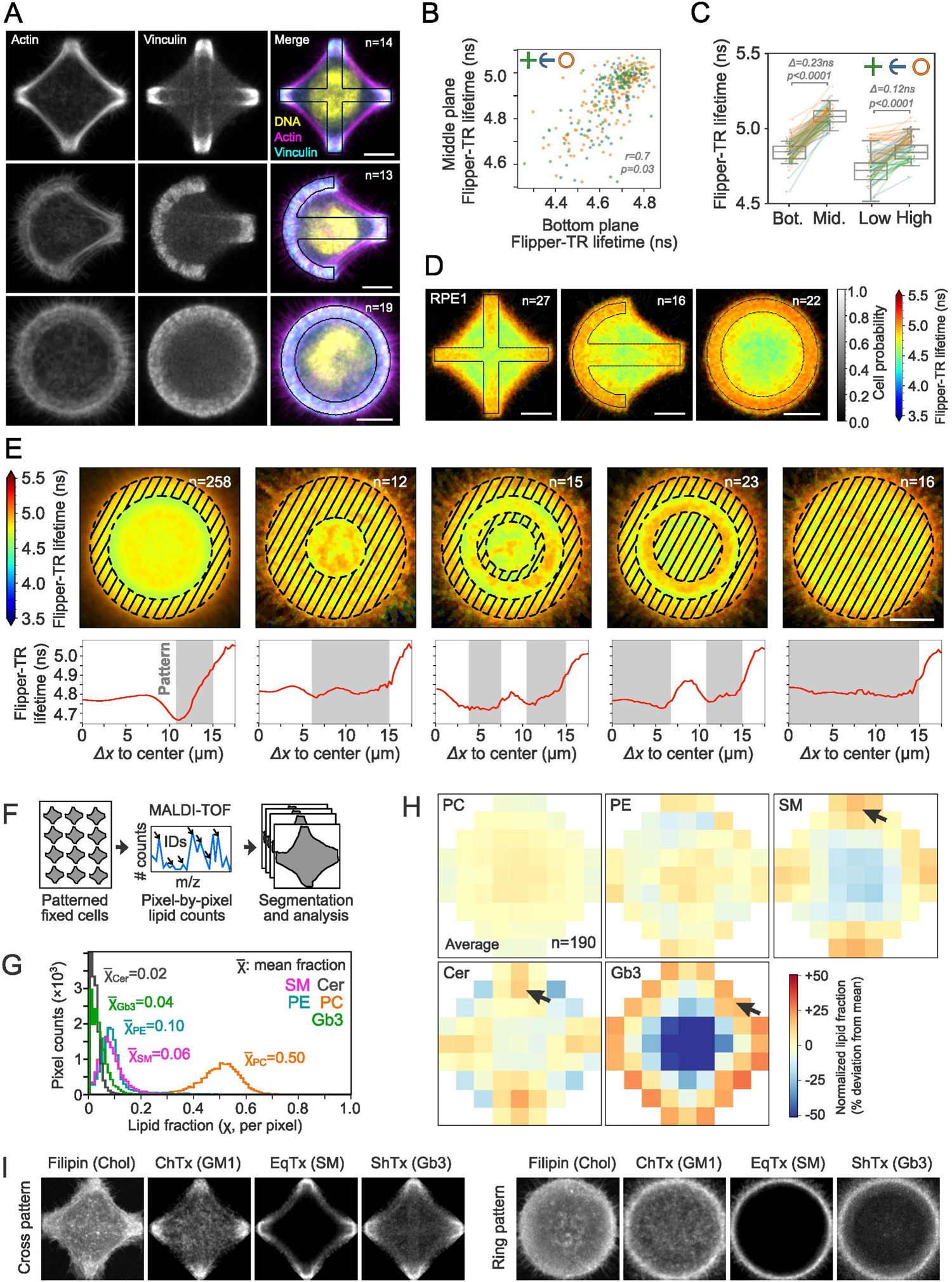
Adhesion and cell morphology shape spatial gradients in membrane tension but not major lipid components. **A**: Average fluorescence images at basal plane of cross (top row, n=14), crossbow (middle row, n=13), and ring (bottom row, n=19) micropatterned cells stained with phalloidin for actin (first column), vinculin antibody (middle column). Last column, color overlay of Hoechst (yellow), vinculin (cyan), and actin (magenta). **B**: Average Flipper-TR fluorescence lifetime (ns) at ventral plane vs. dorsal plane of cross (green), crossbow (blue), and ring (orange) micropatterned cells. Labels refer to r and p values from linear fit. **C**: Average Flipper-TR lifetime (ns) at ventral and dorsal planes (bottom/middle), and bottom and top quintiles (low/high) of cross (green, n=27), crossbow (blue, n=16), and ring (orange, n=22) micropatterned RPE1 cells. Labels represent difference in average values and Welch’s test P value. Data distribution in black. **D**: Cell averaging of average Flipper-TR lifetime in ventral membrane of RPE1 cells on micropatterns. **E**: Top, cell averaging of average Flipper-TR lifetime in ventral membrane of patterned HeLa cells on different motifs. Bottom, average Flipper-TR lifetime vs. distance from pattern center. Shaded areas represent adhesive micropattern location. **F**: Schematics representing MALDI-MSI procedure. **G**: Histogram of pixel-wise lipid fractions, and average lipid fractions for different lipid species: sphingomyelin (SM, magenta), ceramide (Cer, grey), phosphatidylethanolamine (PE, blue), phosphatidylcholine (PC, orange), and globotriaosylceramide (Gb3, green). **H**: Cell averaging of spatial MALDI-MSI of different lipid species. Color scale represents local differences from mean value normalized in terms of standard deviations. Arrows indicate relative lipid accumulation. **I**: Average fluorescence images at basal plane of HeLa cells in cross (left, n=10) and ring (right, n=10) micropatterns stained with filipin, Cholera toxin, Equina toxin, and Shiga toxin targeting cholesterol, GM1, sphingomyelin, and globotriaosylceramide, respectively. **A,D,I**: Scale bar, 10 µm.

**Figure S4:**
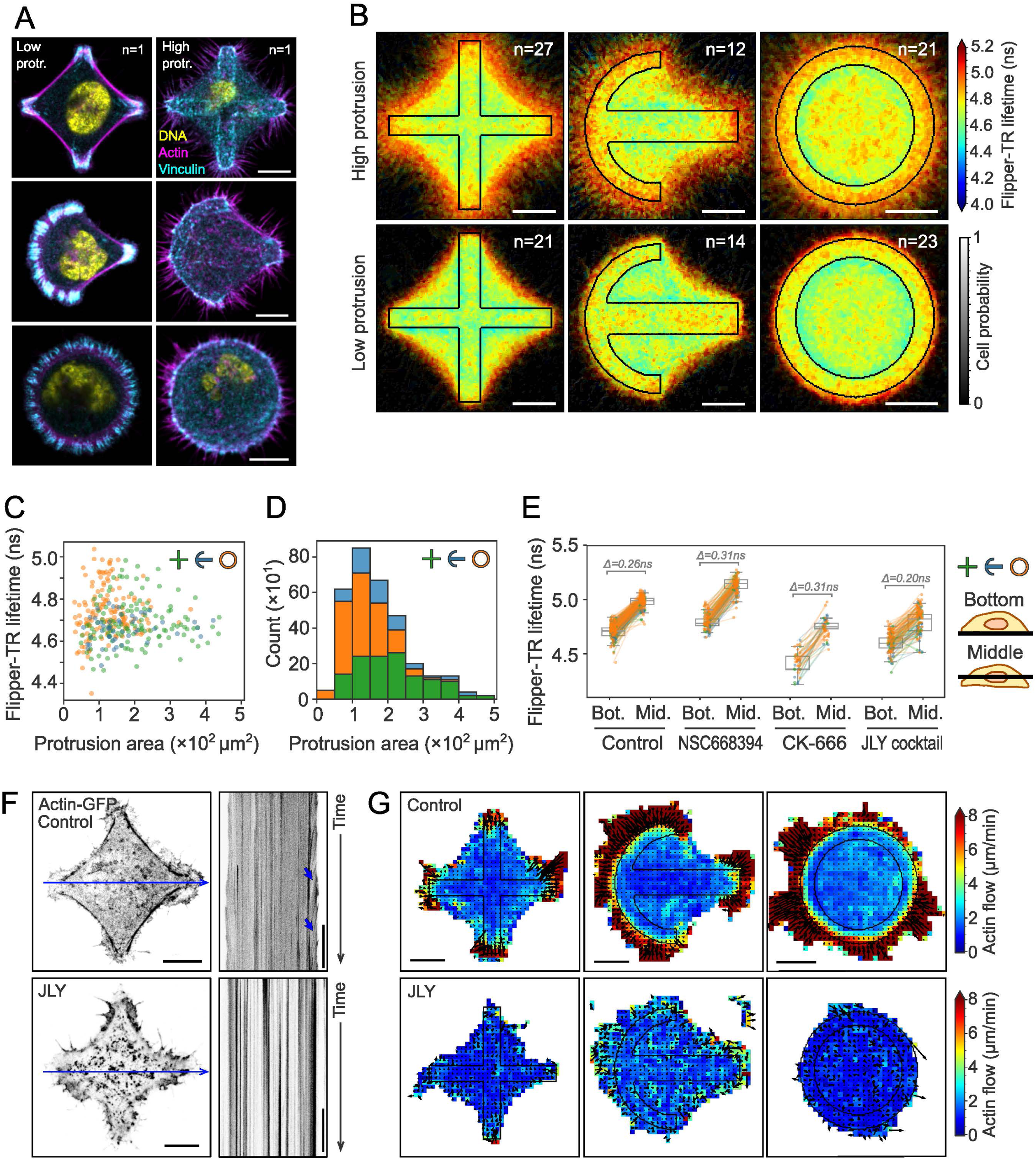
Effect of cell morphology on tension gradients. **A**: Representative fluorescence images at basal plane of cross, crossbow, and ring micropatterned cells stained with phalloidin (magenta), vinculin antibody (cyan), and Hoechst (yellow) showing low (left column) or high number of protrusions (right). **B**: Cell averaging of average Flipper-TR lifetime in ventral membrane of HeLa cells on micropatterns showing high (top) or low (bottom) number of protrusions. **C**: Average Flipper-TR lifetime (ns) per cell at ventral membrane vs. protrusion area (µm²) of cross (green), crossbow (blue), and ring (orange) micropatterned HeLa cells. **D**: Histogram of protrusion areas (µm²) of cross (green), crossbow (blue), and ring (orange) micropatterned HeLa cells. **E**: Average Flipper-TR lifetime (ns) at ventral and dorsal planes (bottom/middle) of cross (green), crossbow (blue), and ring (orange) micropatterned HeLa cells under different drug treatments. Labels display difference from control in ns. Lines represent data pairing. Data distribution in black. **F**: Left, representative images at basal plane of cross micropatterned actin-GFP HeLa cells under control (top) and JLY (bottom) conditions. Blue line indicates kymograph position. Blue arrows mark retrograde flow events. **G**: Cell averaging of particle image velocimetry analysis of actin-GFP HeLa cells on cross, crossbow, and ring micropatterns under control (top, n=10,10,12 respectively) and JLY treatments (bottom n=4,4,4 respectively). Color and arrow lengths indicate cortical flow magnitude. **A,B,F,G**: Scale bar, 10 µm.

**Figure S5:**
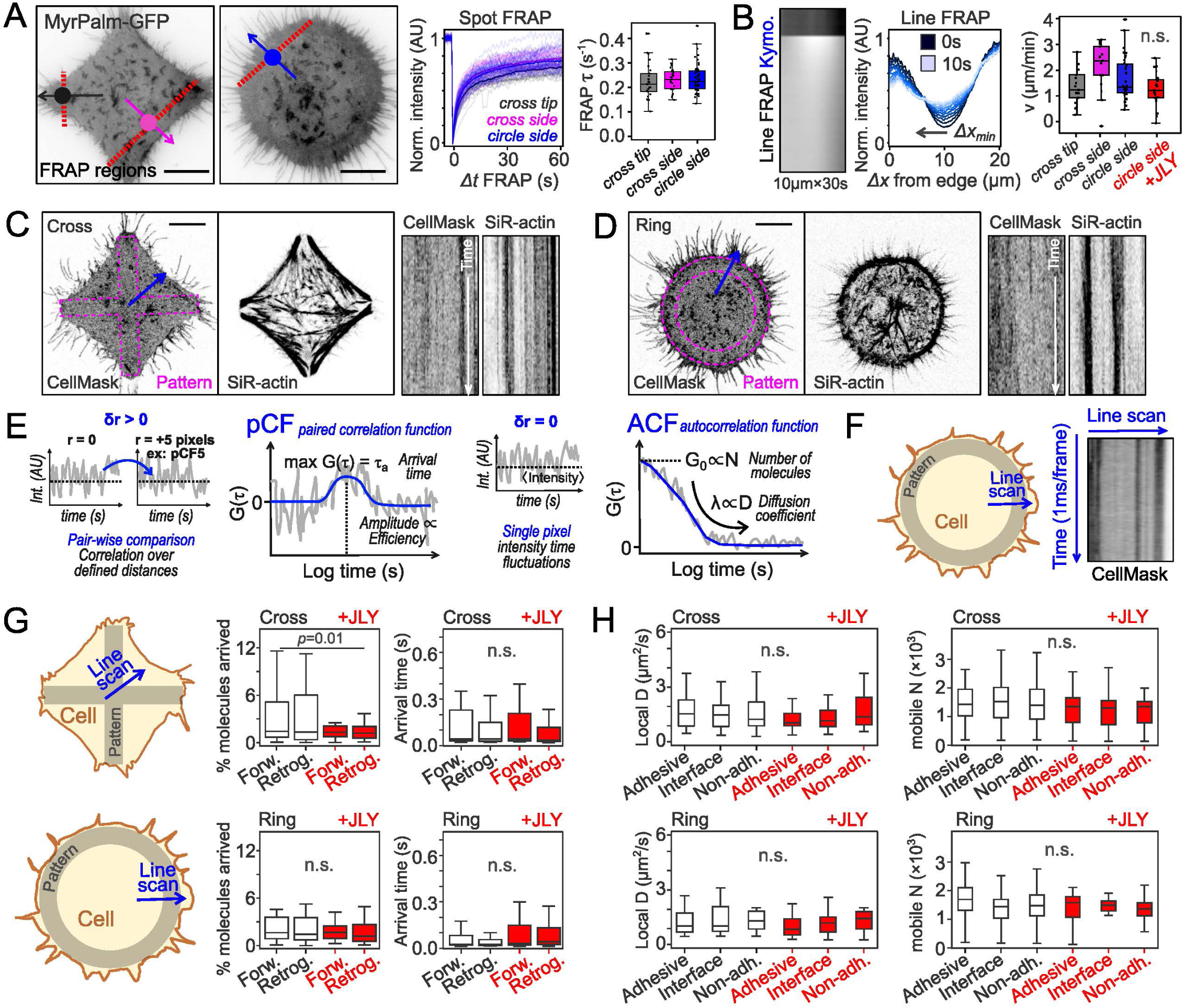
Lipid diffusion and flows are limited in patterned cells. **A:** Left, FRAP experimental design overlaid on images of HeLa MyrPalm-GFP on cross and ring micropatterns. Line FRAP bleaching sites marked with red dotted lines; recovery tracked along black, magenta, and blue arrows. Middle, normalized fluorescence intensity over time after FRAP for spots at cross tip (grey), cross side (magenta), or ring side (blue). Shaded lines represent individual time traces. Right, FRAP recovery constant (s^-1^). **B**: Line FRAP kymograph (inverted LUT) of ring cell from panel A and intensity vs. distance from edge, colored by time elapsed from FRAP. Right, velocity (µm/min) of fluorescence minima vs. distance to cell edge. Statistical test: Welch’s *P* value=0.04. **C-D**: Cell mask and SiR-actin images of HeLa cells on cross (panel C) or ring (panel D) pattern. Kymographs from blue arrows are shown on the right. **E:** Parameters calculated from pair-correlation function (pCF, left) and autocorrelation function fits (ACF, right). **F:** Schematics of the acquisition of line scans, and representative line scan from a cell mask-stained HeLa cell. **G-H**: Schematics of line scan acquisition and fluorescence correlation analysis of patterned HeLa cells stained with CellMask dye under control (black) and JLY (red) conditions. Mean values ± standard deviation. **G**: pCF analysis of cross (top) or ring (bottom) patterned cells. First column, percentage of molecules arrived. Second column, arrival time (s). **H**: ACF analysis of cross (top) or ring (bottom) patterned cells. First column, diffusion coefficient (µm²/s). Second column, number of moving molecules (in thousands). **A-D**: Scale bar, 10 µm

